# Excessive feeding induces development of a colony-like body plan in paratomous flatworms

**DOI:** 10.64898/2026.08.05.742992

**Authors:** Ludwik Gąsiorowski, Monika Sysiak, Denise Klinkenbuß, Katarzyna Tratkiewicz

## Abstract

Colonial animals rely on agametic reproduction to generate repeated colony modules (zooids). Although zooid development has been extensively studied in fully colonial species, such as cnidarians, bryozoans, or tunicates, it provides limited insight into how and why animals transition from solitary asexual fission to colony formation. Here, we fill this gap by studying the dynamics of asexual development in the microscopic flatworm *Stenostomum,* which can alternate between asexual fission and colony-like linear chains, composed of several zooids connected tail-to-head. Combining ecological, developmental, physiological, and transcriptomic analyses, we demonstrate that in four species of *Stenostomum,* chain formation can be triggered by manipulating food availability. This ecological input changes the balance between somatic longitudinal growth and head morphogenesis rate, generating a transient modular organism without requiring a new developmental program. We found no evidence for division of labor among zooids or for enhanced predation avoidance of the worms in chains, indicating that chain formation might be a developmental byproduct without obvious adaptive value. Together, these findings suggest that a neutrally evolving developmental byproduct of food-modulated allometric growth may provide a mechanistic route from asexual reproduction to coloniality.

## Introduction

Emergence of complex life cycles is considered one of the driving forces in evolution, allowing for the rapid appearance of new morphological and developmental characters that significantly increase the overall organismal complexity^1–5^. Among various types of life cycle modifications that evolved in animals, one of particular interest and prevalence is the colonial lifestyle. Animal colonies are composed of modular morphological units (zooids) that are generated asexually through somatic tissues and remain morphologically connected, enabling a mutualized ecology and physiology across the entire colony^2–5^. As such, coloniality strictly depends on the ability for asexual reproduction and enables the evolution of remarkable morphological and ecological diversity, for instance, through labor division and specialization of contributing zooids^3,6–8^. The colonial lifestyle evolved independently in several groups of asexually reproducing animals^2–4^, e.g., in cnidarians, bryozoans, hemichordates, or tunicates, often in concert with the evolutionary success of these clades. Accordingly, the ecological and developmental aspects of coloniality are relatively well studied in those animals^6,9–16^. While studies of fully colonial species can reveal consequences of the evolutionary transition to a colonial lifestyle, they have a limited capacity for elucidating how the animals enter this evolutionary route – i.e., why and how some species gave up individuality and instead evolved towards asexually formed, modularly structured super-organisms. To address these questions, one needs to study animals that are at the edge of what could be defined as a colony.

One such group is Catenulida, a clade of microscopic, free-living, and mostly freshwater flatworms that form a sister group to all the remaining platyhelminths (Fig. 1a, b)^17–20^. These animals predominantly reproduce asexually, through the process known as paratomy^21–26^, in which new posterior and anterior structures develop within the trunk of an adult worm, giving rise to two zooids connected tail-to-head (Fig. 1c). While some catenulids use paratomy only to generate new individuals, others can modify their lifestyle and undergo multiple rounds of paratomous development, without splitting the newly formed zooids^20,27–29^. Then, they form a so-called chain, a modular conglomerate of asexual zooids that remain morphologically connected (Fig. 1c), thus representing a simple colony-like entity. Chain formation seems to be environmentally triggered in some species (e.g., by temperature in *Stenostomum leucops*^29^), while in others it represents a dominant life stage (e.g., *Catenula lemnae*^20,28^), indicating different degrees of incorporation of this colony-like stage into the catenulid life history. Therefore, Catenulida represent an excellent model clade to study how asexual reproduction can be tuned to produce modular, colony-like forms, and to investigate the ecological, physiological, and evolutionary implications of such a developmental shift.

**Fig. 1.**
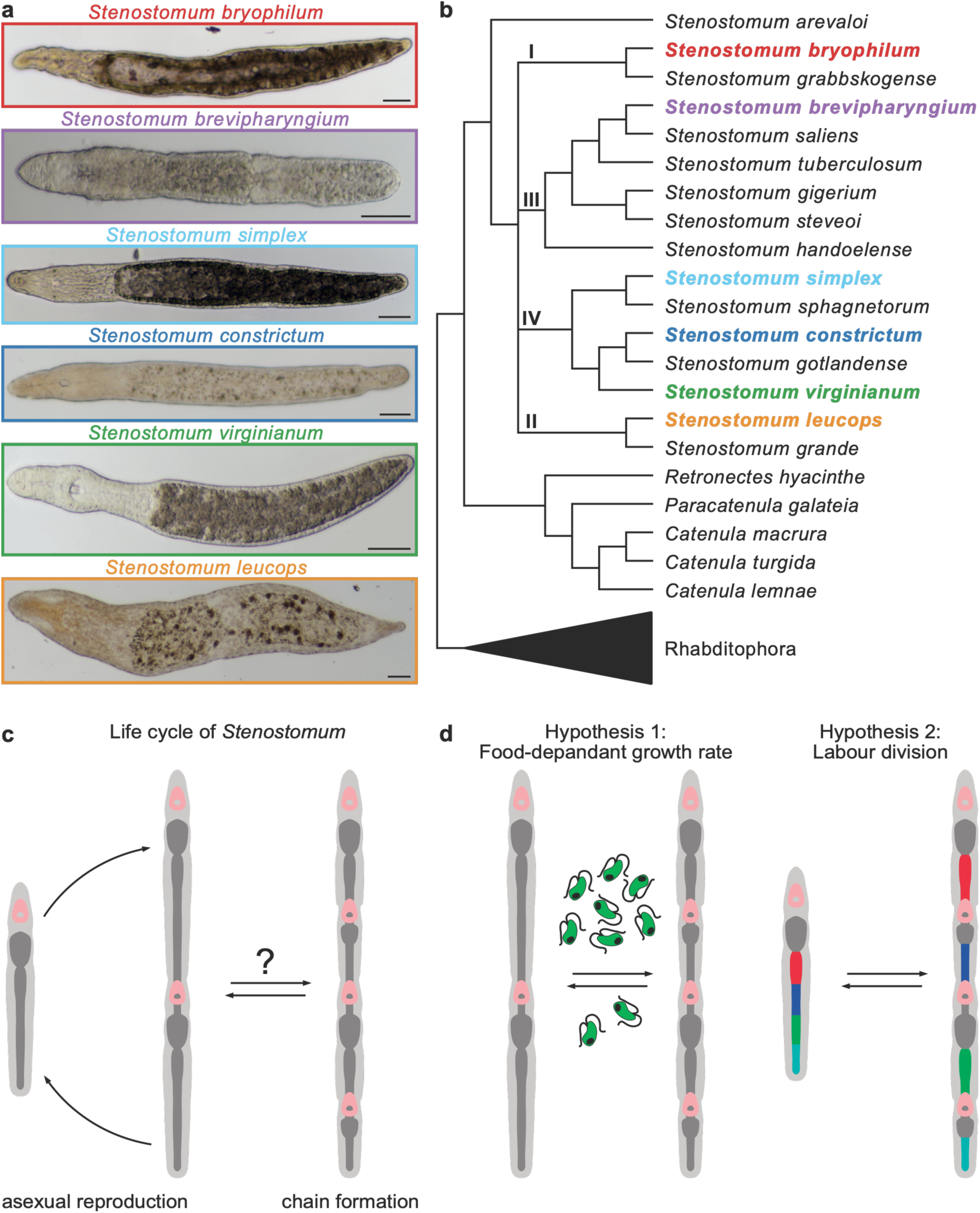
| *Stenostomum* flatworms as a model clade to study chain formation. **a** Microphotographs of six species used in this study; scale bars indicate 50 µm. **b** Position of investigated species in catenulid phylogeny. Major stenostomid clades are indicated with Roman numerals. Tree topology and clade numbering follow the previous study^20^. **c** Alternation between solitary fission and chain formation in the life cycle of *Stenostomum*. **d** Eco-developmental hypotheses tested in this study: 1. chain formation is triggered by excessive food, 2. formation of the chain allows labor division, with particular zooids of the chain playing different roles in the digestive process (colors denote different digestive functions).

To characterize the development and function of these simple colony-like forms, we set out to study details of chain formation in the most speciose catenulid genus, *Stenostomum*, focusing on its ecological context and developmental mechanisms. We wanted to test which factors promote chain formation over solitary fissions and whether the chain formation has any adaptive value. In particular, we wanted to test two eco-developmental hypotheses (Fig. 1d). First, alternation between fission and colony formation might be simply a function of food availability – excessive nutrients accelerate growth, promoting the formation of new zooids. Alternatively, chains might warrant adaptive value, for instance, by allowing labor division, typical for many colonial animals^6–8^. As zooids in chains share a gut, we particularly wanted to test whether modules of the chain perform distinct digestive functions. Finally, we wanted to test whether switching between both modes requires a qualitative shift in developmental control, or whether they represent a quantitative cellular response to changing environmental conditions. By resolving the mechanisms behind alternation of asexual fission and colony formation in catenulids, we hope to provide insight into possible early stages of animal evolution towards coloniality.

## Results and discussion

### Dietary specialization of Stenostomum

First, we tested the dietary specificity of different catenulid species across a range of potential food sources to establish a food source suitable for subsequent feeding experiments. We focused on six species of *Stenostomum*: *S. bryophilum*, *S. brevipharyngium*, *S. simplex*, *S. constrictum*, *S. virginianum*, and *S. leucops,* differing by size and morphology (Fig. 1a) and representing four major clades recently defined within the genus (Fig. 1b)^20^. Catenulid flatworms are considered predators of microinvertebrates and unicellular eukaryotes^20,27,30–38^. We therefore tested seven prey species: the rotifer *Lecane inermis*, the ciliate *Paramecium bursaria*, the cryptophytes *Cryptomonas* sp. and *Chilomonas paramecium*, the euglenid *Euglena gracilis,* and the chlorophytes *Scenedesmus obliquus* and *Chlorella variabilis* (Fig. 2a). These species co-occur with stenostomids in freshwater habitats, varying greatly in size, ecology, cellular metabolism, morphology, and taxonomic affinity (Fig. 2a), thereby constituting a wide and diverse spectrum of potential prey.

**Fig. 2.**
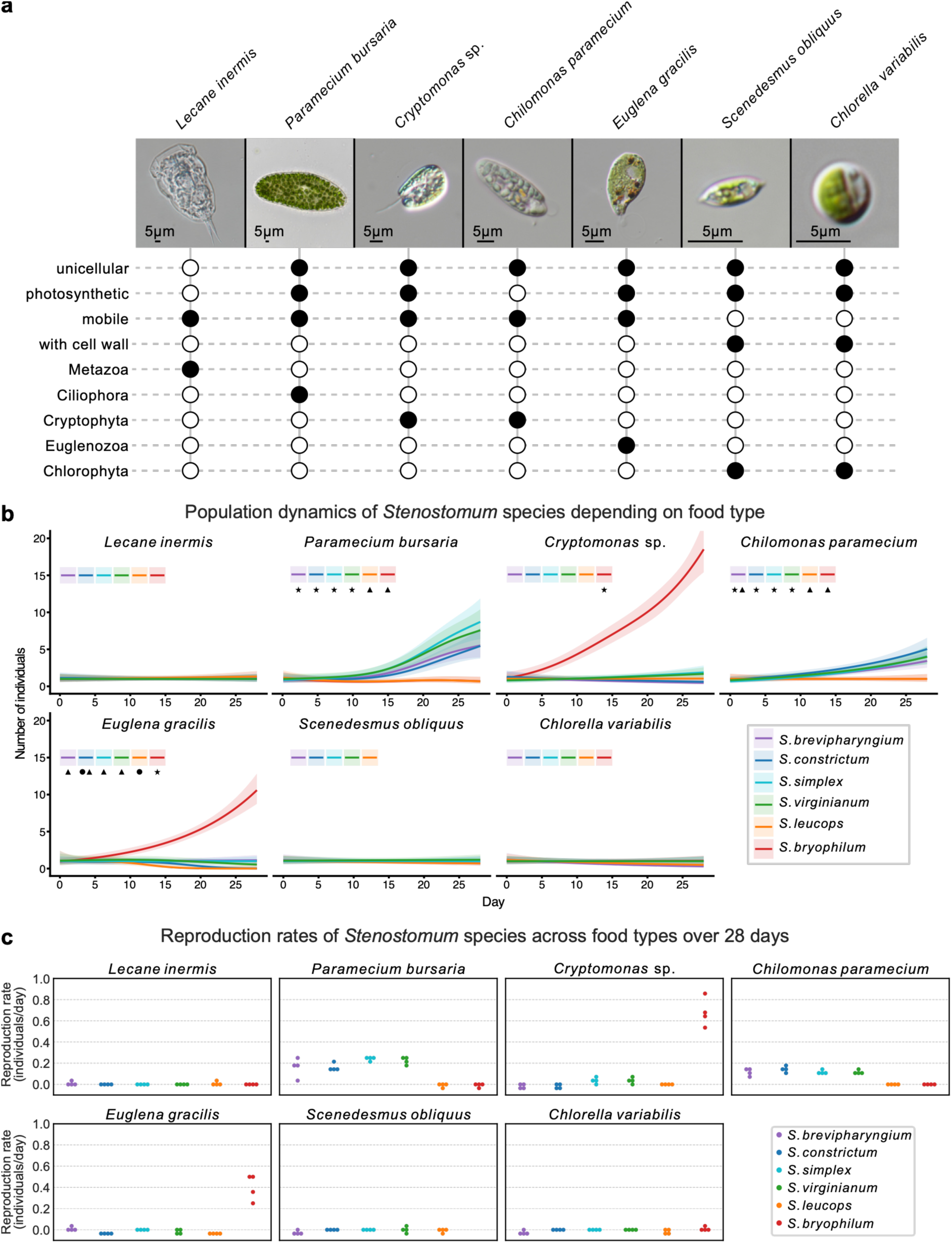
| Dietary specificity of studied *Stenostomum* species. **a** Characteristics of organisms used as food in the feeding experiments. **b** Population dynamics of six *Stenostomum* species fed for 28 days with particular prey items as the sole food source (N = 4 for each species combination). Solid lines depict the population growth trajectories predicted by the generalized additive model (GAM), and lighter areas indicate the 95% confidence intervals. Different symbols (triangles, stars, and circles) indicate significant differences between trajectories. **c** Asexual reproduction rates (RR) (number of new individuals produced over 28 days) of individual founding worms in each experiment. Dots indicate RR for single replication.

Initially, we cultured four individual worms of each catenulid species and fed them *ad libitum* with a single prey species for 28 days. By quantifying population dynamics and reproductive rates for each worm–prey combination, we could detect dietary suitability among *Stenostomum* species (Fig. 2b and c, Supplementary Tables 1, 2). Three food types (rotifer and two chlorophytes) were insufficient to support reproduction in any of the tested *Stenostomum*. *Cryptomonas* sp. and *Euglena gracilis* were efficient food only for *S. bryophilum*, which reproduces rapidly when fed with either organism, but failed to support reproduction in the other species (Fig. 2b and c, Supplementary Tables 3, 4). Interestingly, for most *Stenostomum*-prey interactions, we detected prey cells inside the digestive system of the worms (Extended data Fig. 1). This indicates that insufficient food quality or digestive issues, and not the inability to capture the prey, resulted in the lack of reproduction. Finally, two prey species, *Chilomonas paramecium* and *Paramecium bursaria,* emerged as the most universal food types, allowing reproduction in four out of six tested species, namely *S. brevipharyngium*, *S. simplex*, *S. constrictum*, and *S. virginianum* (Fig. 2b and c, Supplementary Tables 3, 4).

Consistent with previous studies^31,32,35–37^, we found that ciliates represent a preferred food for multiple species of *Stenostomum*, suggesting a strong prey-predator interaction between these two groups. However, our results also showed that *Stenostomum* species display stark differences in dietary specificity. We found that while *S. brevipharyngium*, *S. simplex*, *S. constrictum*, and *S. virginianum* show similar prey preference profiles, *S. leucops* and *S. bryophilum* differ substantially. Particularly, *S. leucops* did not grow on any of the tested food items, whereas *S. bryophilum* was the only species that thrived when fed with *Cryptomonas* sp. and *Euglena gracilis*. Collectively, these results indicate that catenulid species occupy different nutritional niches in their microenvironments, and some species might have evolved as specialized predators of specific unicellular eukaryotes. Notably, feeding with a ciliate *P. bursaria* showed steady and uniform population growth in four tested species of worms, rendering it a universal food type for subsequent experiments.

### Food quantity triggers chain formation

Next, we proceeded to test how the concentration of *P. bursaria* affects the reproduction and development of the worms. First, we exposed different *Stenostomum* species to a known concentration of *P. bursaria* – low (27.5 cells/ml), medium (137.5 cells/ml), and high (275 cells/ml). Individual worms were placed in each food concentration and observed daily for six days. For these observations, we applied pulse feeding, i.e., a single supply of prey at the beginning of the experiment. Reproductive rate of all four species was positively related to food concentration, with limited reproduction at low food concentration and a more pronounced, although not always significantly higher, reproductive response at medium and high food concentrations (Fig. 3a, Extended Data Fig. 2a, Supplementary Tables 5 and 6). Importantly, at the highest food concentration, all species regularly formed chains composed of four to five zooids within approximately 4 days (Fig. 3b).

**Fig. 3.**
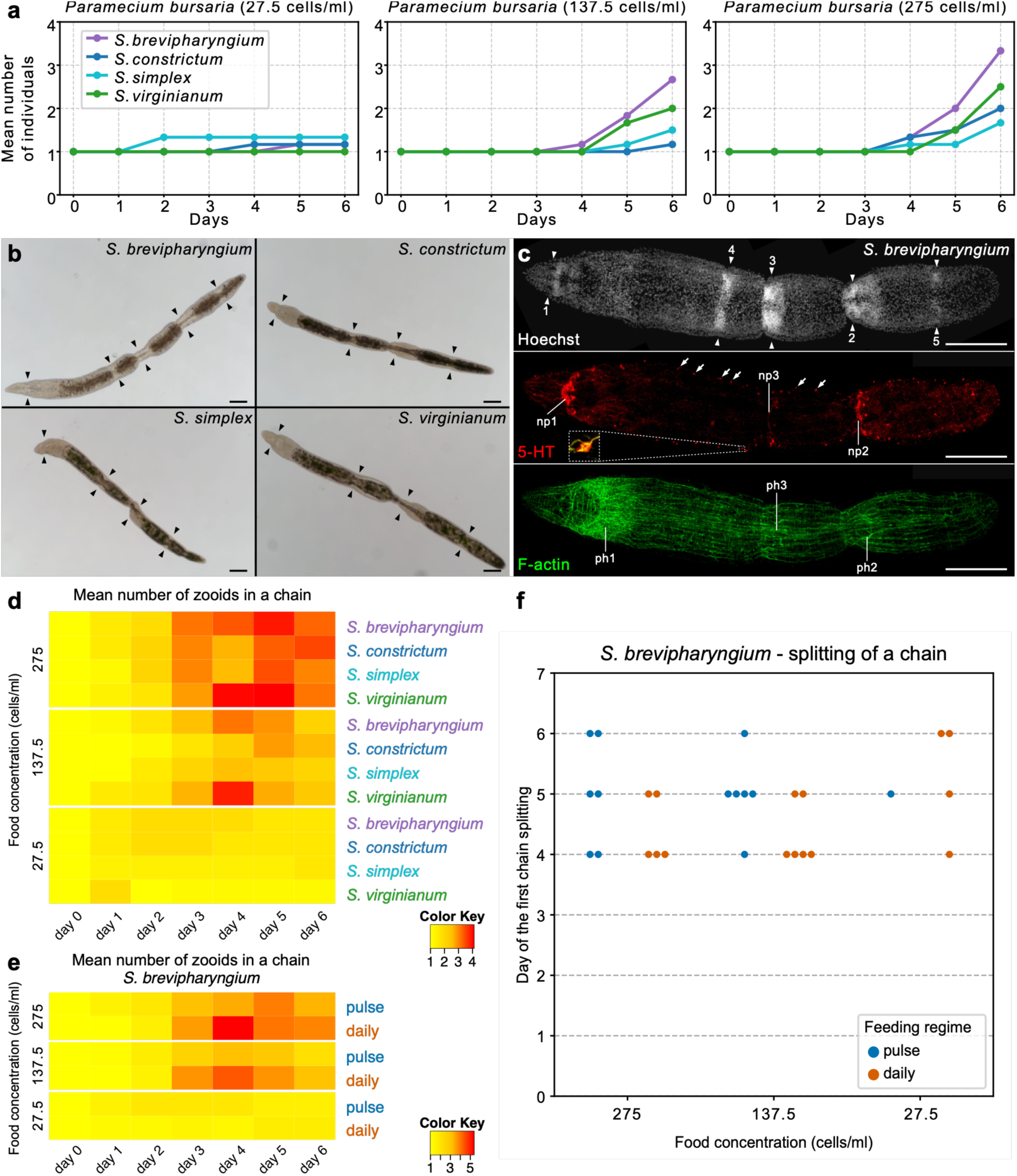
| Chain formation in response to food availability. **a** Changes in mean number of individuals of four species of *Stenostomum* fed with three different concentrations of *Paramecium bursaria* (N = 6 for each species and food concentration combination). **B** Chains formed by all four species in high concentration (275 cells/ml) of *P. bursaria*. Arrowheads indicate head structures. Scale bars represent 100 µm. **c** Detailed organization of a 5-zooid chain of *S. brevipharyngium*. Arrowheads mark head structures at different developmental stages (indicated with numbers: 1 –oldest, 5 –newest). Arrows point to serotonergic perikarya in the lateral nerve cords. Insert shows details of a single perikaryon counterstained against tyrosinated tubulin (yellow). Scale bars represent 100 µm. Abbreviations: *np* brain neuropile, *ph* pharynx. **d** Heatmap depicting changes in the number of zooids per chain in four *Stenostomum* species, fed with different concentrations of *P. bursaria* (N = 6 for each combination of species and food concentration). **e** Heatmap depicting changes in the number of zooids per chain in *S. brevipharyngium* fed with different concentrations of *P. bursaria* under pulse and daily feeding regimes (N = 6 for each combination of species and food concentration). **f** Timing of first chain splitting in *S. brevipharyngium* fed with different concentrations of *P. bursaria* under pulse and daily feeding regimes. The Cox proportional hazards model showed no significant interaction between feeding regime and food concentration (Extended Data Figure 2c).

To investigate the developmental sequence of zooid formation, we studied the detailed morphology of the 5-zooid chains of *S. brevipharyngium*, using fluorescent staining for cell nuclei, nervous system, and musculature (Fig. 3c). Nuclear staining distinguished all five heads, which showed different developmental stages. The oldest new head (head 2) is located in the middle of the chain and has well-developed brain lobes, neuropile, and pharyngeal muscles. The more anteriorly located head 3 possesses some neural and pharyngeal structures but is generally less developed. Heads 4 and 5, located respectively in the anterior and posterior sections of the chain, show the least advanced development and are detectable only as cellular accumulations, typical for early stages of head formation^25,39^. We also observed multiple serotonergic perikarya located along the lateral nerve cords of the 5-zooid chain that are absent in the 2-zooid form of this species^21^. Whether the presence of serotonin in the lateral nerve cords is purely related to feeding status or is involved in the coordination of zooids within the chain remains an open yet intriguing question. Our study of 5-zooid chains proves that new zooids develop in a nested pattern, with new heads inserted into established trunks, resulting in coexistence of modules at different developmental stages.

Next, we quantified how the zooid number changes over time, depending on food concentration (Fig. 3d, Extended Data Fig. 2b, Supplementary Tables 7 and 8). First, we showed that the number of zooids per chain was positively correlated with food concentration. Second, we observed the longest chains around days 4 and 5, regardless of worm species. After this time point, the chains split, giving rise to smaller chains of two to three zooids and reducing the mean number of zooids per chain.

In our pulse feeding experiments, the worms consumed practically all the food by day 4, raising the possibility that the chain splitting resulted from prey depletion. To test this, we compared the mean number of zooids per chain and the timing of chain splitting between *S. brevipharyngium* subjected to pulse or daily feeding (Fig. 3e and f, Extended Data Fig. 2c), with animals in the latter treatment kept in constant food concentration. Although daily-fed animals produced, on average, more zooids per chain, chain fragmentation timing was comparable between treatments (Fig. 3e). Chains consistently split between day 4 and 6, with no significant changes between feeding regimes or food concentration (Fig. 3f, Extended Data Fig. 2c, Supplementary Table 9). *S. brevipharyngium* requires four days for complete head regeneration^21,25,26^, and we infer that the same time is needed for head morphogenesis during paratomy. Consequently, we propose that chains split once the oldest of the newly formed heads is fully developed, suggesting that the duration of head morphogenesis limits the maximum number of zooids per chain.

### Chains are not adaptive

The development of a reliable method to induce chain formation allowed us to test hypotheses about the physiology and ecology of chain-forming *Stenostomum*. First, we tested for the digestive division of labour. A recent single-cell transcriptome of *S. brevipharyngium* identified three cell types with distinct spatial distribution in the digestive system – anterior gut glands (*ST14*^+^), trunk gut glands (*Kazal1*^+^), and hindgut cells (SLC26A2^+^)^26^. These cell types are strongly segregated into the anterior, middle, and posterior gut in single-zooid animals (Fig. 4a) and likely have specialized functions in food processing. Digestive division of labour would predict an uneven distribution of digestive cell types among zooids (Fig. 1d). However, *in situ* hybridization of the respective marker genes for these cell types in the 4-zooid chains of *S. brevipharyngium* (Fig. 4a) showed their repetitive distribution within each zooid. Only the anterior gut glands showed postponed development, being absent in the most recently emerged zooids, consistent with delayed development of the anterior structures during chain formation. Overall, these expression patterns indicate that each zooid has digestive capacities comparable to a solitary worm, with no spatial segregation of digestive cell types.

**Fig. 4.**
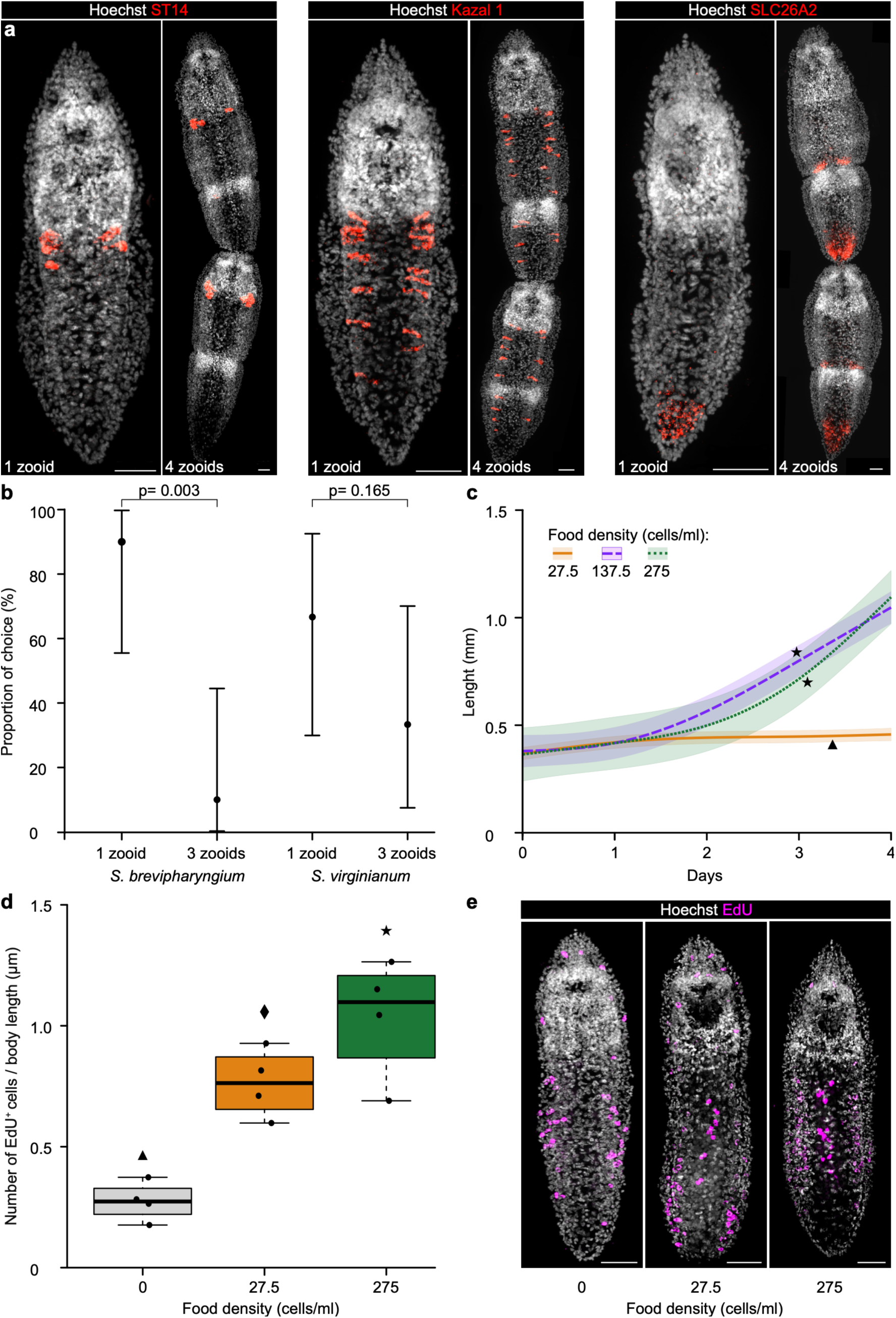
| Impact of the feeding status on the biology of the worm. **a** RNA *in situ* hybridization chain reaction of molecular markers for digestive cell types allows comparison of their distribution between single worms and 4-zooid chains. The name of the hybridized gene is provided in each panel; scale bars represent 20 µm. **b** Predator preference (%) between 1-zooid and 3-zooid prey within the two prey species *S. brevipharyngium* (N = 10) and *S. virginianum* (N = 9) predated by *S. sphagnetorum*. Points represent the estimated model probabilities of prey choice, and the vertical error bars indicate the 95% confidence intervals. p-values denote the comparisons between prey sizes within each species. **c** Longitudinal growth rate of *S. brevipharyngium* exposed to different concentrations of *P. bursaria* (N = 8 for each condition). Solid lines depict growth trajectories of body length predicted by the generalized additive mixed model (GAMM), and lighter areas indicate the 95% confidence intervals. Different symbols (triangles and stars) indicate significant differences between trajectories. **d** Differences in the number of EdU^+^ cells adjusted for body length in *S. brevipharyngium* worms exposed to different feeding conditions (N = 4 for each condition). Boxes represent the interquartile range, horizontal lines indicate medians, and whiskers show the range of observed values. Different symbols (triangle, diamond, and star) indicate significant differences between food concentrations. **e** The visualization of EdU^+^ cells in *S. brevipharyngium* exposed to different feeding conditions. Scale bars represent 20 µm.

Many aquatic invertebrates increase their body size as a mechanism of predator avoidance^40–42^. Since *Stenostomum* chains are considerably longer than single worms, we tested whether chain formation reduces predation. We exposed single and 3-zooid individuals to a predator for 48 h and compared their survival. For each experiment, we used *S. brevipharyngium* or *S. virginianum* as prey, and *Stenostomum sphagnetorum,* a facultative predator of other stenostomids. For *S. brevipharyngium*, the 3-zooid chains were significantly less likely to be chosen as prey, whereas survival did not differ between 1– and 3-zooid individuals of *S. virginianum* (Fig. 4b, Supplementary Table 10). Therefore, chain formation might protect from predation in some instances, but it does not represent a universal mechanism of predator avoidance.

We tested two potential adaptive benefits of chain formation – digestion-oriented labor division of zooids and predation avoidance – and found no convincing evidence for either. Consequently, we suggest that chain formation is not necessarily adaptive. Instead, it might have evolved as a neutral character mechanistically related to paratomous development.

### Feeding affects the somatic growth rate

To explain how chain formation can neutrally originate from paratomy, one needs to understand the dynamics of paratomous development. Production of new individuals in the context of paratomy is a combined result of two parallel processes – morphogenesis and somatic growth. Earlier, we showed that the head morphogenesis rate is constant and independent of feeding status. Now, we wanted to test how food availability affects somatic growth.

First, we measured body length changes of *S. brevipharyngium* fed with different concentrations of *P. bursaria* over 4 days (Fig. 4c). While there was no difference in longitudinal growth rates between the medium and high food conditions, both differed significantly from animals fed with low food concentration (Supplementary Tables 11 and 12). As growth in *Stenostomum* relies on stem cell division^23,26,43^, mitotic activity provides another proxy for somatic growth rate. Therefore, we measured the number of actively dividing cells in *S. brevipharyngium* worms that were either starved or fed with low or high food concentrations. To assess cell division rates, animals were kept for 24 h in each condition and pulsed for 3 h with 5-Ethynyl-2′-deoxyuridine (EdU), which incorporates into the mitotically active cells. We found that the number of EdU^+^ cells, normalized to animal length, increased significantly with the feeding status (Fig. 4d and e, Supplementary Tables 13 and 14). Altogether, our results show that somatic growth in *S. brevipharyngium* depends on food availability, with higher food concentrations enhancing longitudinal growth.

### Transcriptional response to increased food uptake

Next, we complemented ecological and physiological analyses with transcriptomics to examine systemic responses to increased food uptake, which initiates the process of chain formation. To gain insight into gene expression changes evoked by different feeding conditions, we sequenced RNA from *S. brevip*haryngium worms subjected for 48 h to low or high concentrations of *P. bursaria* (three biological replicates of 60 worms per condition). Differential gene expression analysis identified genes upregulated in animals fed with low or high food concentrations (Fig. 5a, b). Altogether, we identified 1241 genes significantly upregulated in either condition (|log_2_fold change| > 2, padj < 0.05), of which 42.2% were upregulated under high food and 57.8% under low food conditions (Fig. 5b). To determine cell-type specific responses to changes in food availability, we examined cell-type specificity of the differentially expressed genes using the single-cell atlas of *S. brevipharyngium*^26^. 402 out of 1241 differentially expressed genes were specific to at least one cell cluster, of these 121 (30%) were upregulated in high food and 281 (70%) in low food conditions. Genes upregulated in animals exposed to a high food concentration were mostly specific to intestinal cell types and stem cells (Fig. 5c). We suggest that while the response in intestinal cells is directly related to the digestion process, the activation of stem-cell-specific genes correlates with increased somatic growth in animals exposed to high food concentration. Conversely, genes upregulated in animals exposed to a low food concentration were specific to multiple cell types, including intestinal cells, epidermis, neurons, and muscle. Interestingly, many genes upregulated in lowly-fed animals were specific to ciliated receptors (77 genes), a specialized cell type present in the epidermis and pharyngeal lining of *S. brevipharyngium*^26^. While the exact function of these cells remains unknown, they might be involved in prey sensing or mucus production, which is known to facilitate predation in *Stenostomum*^27,38^. Therefore, we suggest that the upregulation of genes specific to ciliated receptors in lowly-fed animals might reflect their active hunting state. Overall, these data indicate that specific cell types exhibit distinct responses to feeding status in *S. brevipharyngium*, likely reflecting both feeding physiology and the initiation of developmental responses.

**Fig. 5.**
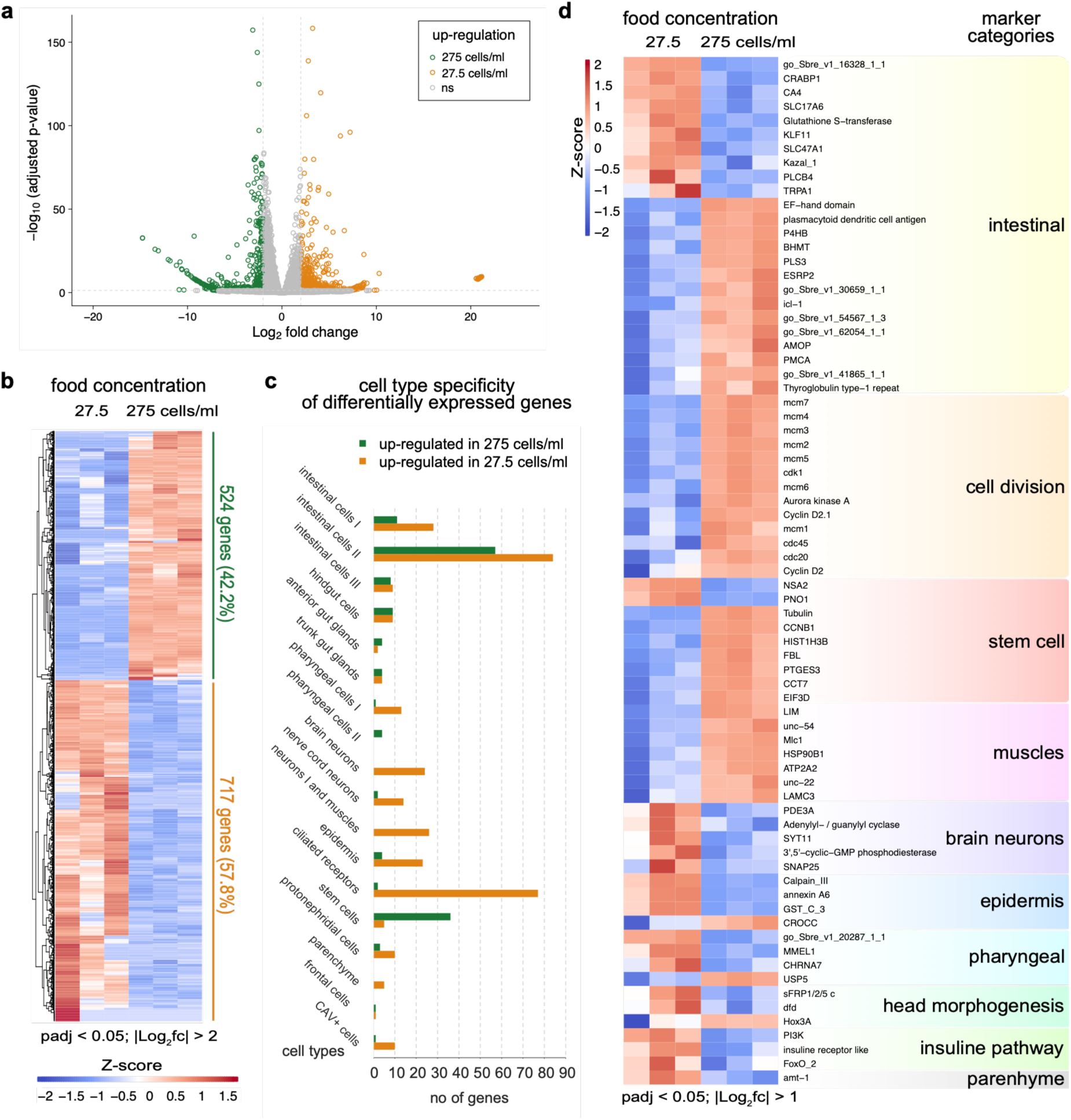
| Effects of food concentration on gene expression and cell-type-specific transcriptional responses in *Stenostomum brevipharyngium*. **a** Volcano plot showing significance (adjusted *p* < 0.05, two-sided Wald test) and magnitude of change (|log_2_FC| > 2) for differentially expressed genes (DEGs) in response to low (27.5 cells/ml) and high (275 cells/ml) food conditions. **b** Heatmap of normalized expression (row Z-scores) of DEGs (adjusted *p* < 0.05, two-sided Wald test; |log_2_FC| > 2). **c** Comparison of significantly upregulated DEGs between food conditions across cell-type clusters. Enrichment was defined as higher expression in a given cluster compared to all others (two-sided Wilcoxon rank-sum test, adjusted *p* < 0.05; |log_2_FC| > 1). **d** Heatmap showing expression changes of different categories of marker genes in response to food concentration (adjusted *p* < 0.05, two-sided Wald test; |log_2_FC| > 1).

Next, we investigated the expression dynamics of a set of known marker genes (Fig. 5d), including various cell type markers and genes involved in cell division, head morphogenesis, and the insulin signaling pathway, a major regulator of nutritional responses in many animals^44,45^. For some of these categories, we observed condition-specific upregulation (e.g., cell division markers), while others show bidirectional expression shifts (e.g., intestinal markers). Genes involved in cell division and most of the stem cell markers show upregulation in high food concentration, aligning with stimulated somatic growth in such conditions. Muscle markers were also clearly upregulated in high food conditions, possibly reflecting trunk muscle extension during longitudinal growth. Head-associated genes – i.e., markers of brain neurons and pharynx, and genes involved in head morphogenesis – are upregulated in low food concentration, consistent with cell-type specificity analysis. This can be explained by the fact that food consumption stimulates allometric growth, with trunk tissues growing faster than head structures, resulting in a smaller proportion of head tissues in larger, well-fed animals. A similar developmental allometry has been recently reported in another group of flatworms, planarians, in which larger individuals contain a proportionally smaller share of head tissues^46^. Intestinal markers show bidirectional responses, with 10 markers upregulated in high-food conditions and 14 in low-fed animals, mirroring the cell-type specificity analysis. As genes expressed in the digestive system play diverse, sometimes opposite, roles in food processing, nutrient absorption, and storage, we suggest that this bidirectional response is a consequence of the complex functional landscape of intestine-specific genes. Finally, only three out of 26 tested components of the insulin signaling pathway (*PI3K*, *insulin receptor-like*, and *foxO2*) showed significant differences in expression levels between the high– and low-food conditions, with all three being upregulated in well-fed animals. Therefore, we could not confirm that the insulin pathway plays an important role in the processing of feeding status information in *S. brevipharyngium*.

Altogether, our transcriptomic data show that feeding status has a profound impact on gene expression profiles in *S. brevipharyngium*. Specifically, we observed transcriptional imprints of both feeding physiology and growth mode. The latter is particularly important for understanding the chain formation process and can be characterized by food-induced activation of stem cell proliferation, resulting in allometric development, in which trunk tissues grow faster than head-associated organs.

## Conclusions

Food availability and feeding status can have a prominent impact on developmental processes^44^, as shown in the post-embryonic development of cnidarian tentacles^45^ or the maturation of annelid larvae^47^. In this study, we demonstrated that food availability alone is sufficient to modulate the life cycle of a microscopic flatworm *Stenostomum*, providing yet another example of the direct dietary impact on the developmental process. We propose a simple eco-developmental model that may explain how chain formation can be triggered by food availability (Fig. 6). Asexual development of *Stenostomum* is controlled by two factors: longitudinal growth and head morphogenesis rate. The former allows reaching the critical body length at which the formation of new structures is initiated in the middle of the trunk. The rate of head morphogenesis, on the other hand, dictates the timing of the chain splitting. We showed that while the longitudinal growth rate is food-dependent, the head morphogenesis rate does not follow the same logic and is rather fixed. This results in allometric growth in which, in response to abundant food, trunk tissues grow faster than new heads are being formed. Consequently, in high food conditions, the trunk reaches the length required to initiate a new round of paratomy before head tissues are fully formed, triggering the formation of a chain, whose length depends directly on food availability. However, these chains are not permanent structures and start to fall apart as soon as the oldest of the new heads is fully formed. Importantly, according to this model, paratomy and chain formation are not two different developmental modes. Instead, they represent two outcomes of the same developmental program, determined solely by nutritional input to the system.

**Fig. 6.**
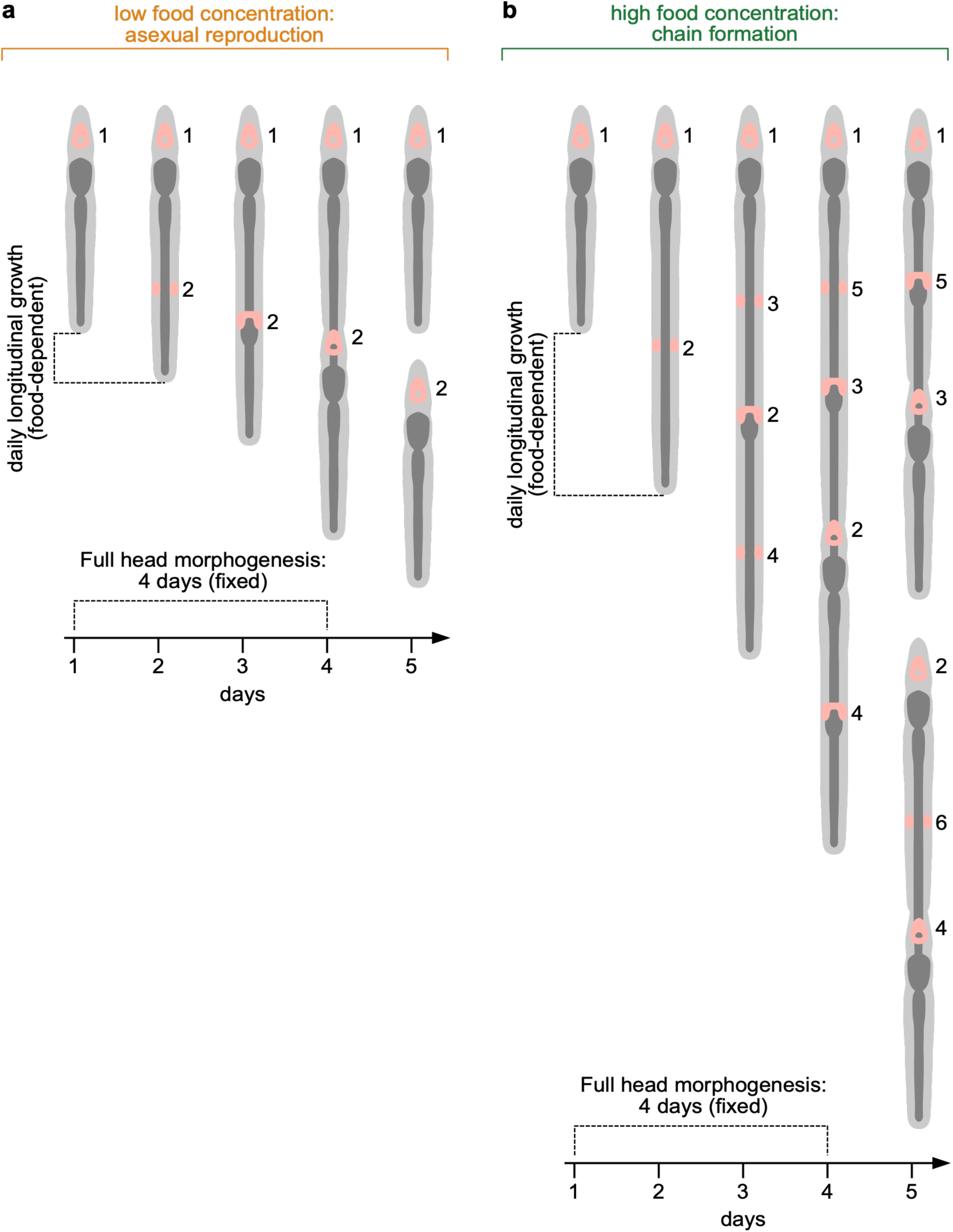
| Eco-development model for food-driven mechanism of alternation between asexual reproduction and chain formation in *Stenostomum*. Asexual reproduction is governed by two parallel processes: longitudinal growth (food-dependent) and head morphogenesis (food-independent). In low food conditions, new heads fully form before the trunks of the progeny reach the length required for paratomy initiation. In high food conditions, trunk tissues grow faster than the new heads are formed, allowing initiation of new paratomous growth zones nested within the trunks of underdeveloped zooids. This gives rise to a chain that splits into smaller ones as soon as the oldest newly formed head is fully developed. Numbers indicate the order in which new heads develop in each condition. Brain tissues are depicted in pink, and the digestive system (with pharynx) in dark grey.

We showed that chain formation does not necessarily bring any obvious benefits to the animal and can hardly be interpreted as an adaptation. Instead, we argue that chain formation should be considered a developmental byproduct of allometric growth and length-dependent onset of paratomy. Such neutrally evolving traits, derived from physical or topological constraints on developmental processes, are often referred to as spandrels^48^. Spandrels may, in some circumstances, acquire adaptive functions (the so-called “exaptations”^49^), and here we suggest that chain formation represents a preadaptation to the colonial lifestyle. Although chains of *Stenostomum* share key features with colonial organisms, including modular organization and asexual formation, they remain transient and do not allow division of labor among zooids. For those reasons, we propose referring to them as paracolonies. However, within catenulids, paracolonial *Stenostomum* coexist with the genus *Catenula*, in which even fully formed zooids do not separate. Instead, chains of some species are composed of numerous (>15) permanently connected zooids, with fully functional heads^20,28^. This indicates that catenulid paracolonies may give rise to a more advanced colonial integration, in which fully developed individuals do not separate from the chain and function as a permanent super-organism. Therefore, we propose that catenulids represent a case in which true coloniality evolved from a developmental spandrel of asexual reproduction modulated by food availability. A similar mechanism may lie at the dawn of the colonial lifestyles of the other animals and represent a universal route from agametic asexual propagation to true coloniality.

## Methods

### Collection and husbandry of animals

The animals used in the experiments came from the collection of the Comparative Invertebrate Zoology Group at the University of Warsaw: *S. brevipharyngium* (UW-0001) was collected in the USA; *S. constrictum* (UW-0002) in the UK; *S. leucops* (UW-0014) in Italy; while *S. simplex* (UW-0009), *S. virginianum* (UW-0016), and *S. bryophilum* (UW-0036) were sampled in Poland. All those species are freshwater organisms and occur in small, temporary water bodies such as peat bogs, ponds or ditches. Species were identified based on morphological characters and validated with multiple molecular barcodes^20^. Animal cultures were started from single individuals (isolines) and kept in 9 cm plastic Petri dishes with Chalkley’s Medium (CM: 1.71 mM NaCl, 0.05 mM KCl, 0.05 mM CaCl_2_) at 20 °C in the dark. Cultures were supplied with wheat grain infusions, which support the growth of a multitude of unicellular organisms serving as food.

### Feeding experiments

All feeding experiments were performed in individual wells of 24-well plates in CM at 20 °C in the dark. To define the food preferences of each species (*S. brevipharyngium*, *S. constrictum*, *S. leucops*, *S. simplex*, *S. virginianum* and *S. bryophilum*) and assess the impact of food type on population growth, worms were starved for 5 days and then individually put in wells containing 1 mL of medium with different types of prey (rotifer *Lecane inermis*, ciliate *Paramecium bursaria*, cryptophytes *Cryptomonas* sp. and *Chilomonas paramecium*, euglenid *Euglena gracilis*, and chlorophytes *Chlorella variabilis* and *Scenedesmus obliquus*). If the prey was originally cultured in a medium other than CM, it was concentrated and resuspended in CM. For each combination of worm and prey species, we used four replicates. The experiment was carried out for 28 days, and the number of animals in each well was counted on Mondays, Wednesdays, and Fridays. Additionally, every Friday, animals were fed if the food was decreasing in a specific well.

To measure the impact of *P. bursaria* concentration on chain formation, first, a dense culture of *P. bursaria* was resuspended in CM. Then the number of cells was counted under the dissecting microscope (in 20 drops, each 20μl), and the suspension was diluted with CM to a working concentration of 275 cells/mL, and then further to 137.5 cells/mL (1:1) and 27.5 cells/mL (1:9). The animals (*S. brevipharyngium*, *S. constrictum*, *S. simplex*, *S. virginianum*) were starved for 3 days and put individually in wells containing 0.5 mL of *P. bursaria* suspension at one of three concentrations (pulse feeding), with six replicates for every species– concentration combination. The worms were counted every day for a week; additionally, the number of zooids in each individual and the day of first chain splitting were noted.

To investigate the impact of constant food concentration on the chain stability of *S. brevipharyngium,* we used a daily feeding regime. The animals were starved for 3 days and then placed in wells with 0.5 mL of *P. bursaria* suspension at three different concentrations: 275, 137.5, and 27.5 cells/ml. The total number of individuals and the number of zooids were counted every day for a week. After counting, 440 μL of medium was removed, leaving 60 μL containing no *P. bursaria* cells, and replaced with the same amount of fresh medium with adequate *P. bursaria* concentration. Additionally, for each experimental well, the day of the first chain splitting was noted.

### Longitudinal growth rate

1-zooid individuals of *S. brevipharyngium* were starved for 3 days, placed individually in 24-well plates (ten replicates for each condition), and fed once with one of three different concentrations of *P. bursaria*: 275, 137.5, and 27.5 cells/ml in CM. Body length of each individual was measured daily for five consecutive days directly in the well using a dissecting microscope Leica Investa-3 with integrated camera.

### Predation avoidance experiment

To test the effect of prey size (number of zooids) on the predator choice, we used 1-zooid individuals of the predatory flatworm *S. sphagnetorum*, starved for 24 h before the experiment. Based on our observations, this period was sufficient to stimulate hunting activity, whereas longer starvation negatively affected the condition of this species. As prey, we used 1-zooid and 3-zooid individuals of either *S. brevipharyngium* or *S. virginianum.* 3-zooid prey were obtained by placing 1-zooid individuals in Petri dishes with ad libitum *P. bursaria* three days before the experiment. To standardize hunger state, 1-zooid prey were also kept for 24 h in dishes with highly concentrated *P. bursaria*, this period was sufficient for food acquisition but too short to produce new zooids. Two hours before exposure to the predator, all prey were transferred to CM to clear their guts. For each replication, one predator was placed in a well containing 0.5 mL CM together with one 1-zooid and one 3-zooid prey. The experimental wells were kept at 20 °C in darkness. We recorded which prey size was consumed first by observing experimental wells at 24 h and 48 h.

### Antibody staining

The animals were first anesthetized in 1.44% (mass:volume) MgCl_2_ in CM for ca. 10 min and then fixed in 4% formaldehyde in Phosphate-Buffered Saline buffer with 0.1% Tween-20 (PTw) for 30 minutes at room temperature (RT). The fixed animals were rinsed three times in PTw, then three times in PBS + 0.1% bovine serum albumin + 0.1% Triton X (PBT) for 15 minutes at RT and afterwards incubated for 30 minutes in 5% normal goat serum dissolved in PBT (PBT+NGS), also at RT. Next, we incubated worms overnight at 4 °C in primary antibodies dissolved in PBT+NGS. We used the following primary antibodies: mouse anti-tyrosinated tubulin, Sigma T9028 (dissolved at 1:500); and rabbit anti-serotonin (5HT), Sigma S5545 (dissolved at 1:250). The next day, worms were washed six times with PBT for 15 min and incubated for 30 min in PBT+NGS, both at RT. The animals were subsequently incubated overnight at 4 °C in secondary antibodies dissolved in PBT+NGS. We used the following secondary antibodies: goat anti-mouse, conjugated with Alexafluor488, Thermo Fisher A-11001; and goat anti-rabbit, conjugated with Alexafluor647, Thermo Fisher A-21244 (both at a concentration of 1:250). After incubation in secondary antibodies, all steps were performed at RT. The animals were washed three times in PBT, two times in PTw, and incubated for 40 min in Hoechst 33342 dissolved in PTw (1:5000). Then, we rinsed the worms twice in PBT and incubated for 1 h in Phalloidin, conjugated with Alexafluor555, Thermo Fisher A34055 (10U/ml dissolved in PBT). Finally, animals were mounted in Fluoromount G (Thermo Fischer, 00-4958-02) and stored at 4 °C for tissue clearing and hardening of the mounting medium.

### In situ RNA hybridization

For fluorescent RNA *in situ* hybridization chain reaction (HCR) v3.0^50^, we used the same DNA probe oligo pools as in the previous study on cell types in *Stenostomum*^26^, ordered at Integrated DNA Technologies. For *in situ* staining, the worms were first anesthetized in 1.44% (mass:volume) MgCl_2_ in CM for ca. 10 min and fixed in 4% paraformaldehyde dissolved in PTw for 30 min at RT. Then animals were washed six times with PTw, transferred to 100% methanol (Me-OH), and stored at –20 °C overnight. The next day, the material was rehydrated in a Me-OH/PTw series at RT, washed four times in PTw, and prehybridized for 40 min at 37 °C in hybridization buffer (30% formamide, 5x SSC, 9mM citric acid pH=6.0, 0.1% Tween-20, 50μ/mL heparin, 1x Denhardt’s solution, 10% dextran sulfate). Afterwards, the animals were placed in HCR probe mixtures at 1μM concentration in hybridization buffer and incubated overnight at 37 °C. The next day, probes were removed with four washes (each 10 min at 37 °C) of probe wash solution (30% formamide, 5x SSC, 9mM citric acid pH=6.0, 0.1% Tween-20, 50μg/mL heparin), followed by three washes in 5xSSC+0.1% Tween-20 at RT. Then, the worms were incubated for 30 min at RT in amplification buffer (5x SSC, 0.1% Tween-20, 10% dextran sulfate). In the meantime, HCR hairpins (Molecular Instruments: B1H1-546, B1H2-546) were prepared by heating for 1 min 30 sec at 95 °C and cooling at RT in the dark for 30 min. The hairpins were mixed in amplification buffer to a final concentration of 40nM and added to the samples. The animals were incubated in the amplification buffer with hairpins overnight at RT in the darkness. The next day, the samples were rinsed three times with 5xSSC+0.1% Tween-20, twice with PTw, and incubated for 40 min in Hoechst 33342 in PTw (1:5000). The stained specimens were washed once in PTw, mounted in Fluoromount G (Thermo Fischer, 00-4958-02), and stored at 4 °C.

### EdU staining

For EdU incorporation experiments, we transferred worms into clean CM and starved them for two days. After that period, worms were moved individually into wells with appropriate food concentration of *Paramecium bursaria* suspended in CM (high food: 275 cells/mL, low food: 27.5 cells/mL; four replicates per food concentration) or to pure CM (starved animals) and left for 24 h. The next day, worms were transferred to fresh, clean CM for 3 h and then soaked for 3 h in 400 μM EdU in CM with the addition of 3% DMSO. For EdU detection, we used the Click-iT™ EdU Cell Proliferation Kit for Imaging, Alexafluor647 (Thermo Fischer, C10340), according to the manufacturer’s recommendations with the following modifications: after incubation, worms were washed several times in fresh CM, anesthetized with 1.44% MgCl_2_ in CM (mass:volume), and fixed with 4% formaldehyde in PTw for 1 h. In addition, incubation in the Click-iT® reaction cocktail was extended to 1 h at room temperature.

### Imaging

Daily observation and length measurements were performed with the Leica dissecting microscope Investa 3 with an integrated camera. Microphotographs of live animals were taken on a Nikon Eclipse NI-SSR compound microscope with a Nikon DS-Ri2 camera. Fluorescently labelled samples were imaged on an Inverted Zeiss Axio Observer Z.1 equipped with a CSU-X1 spinning disc unit and 2x sCMOS Camera Prime BSI (Teledyne Photometrics). The microphotographs and Z-stacks were adjusted for brightness and contrast and further processed in the image analysis software Fiji^51^.

### RNA sequencing and transcriptomic analyses

The worms were kept in CM with 275 cells/ml (high food concentration) or 27.5 cells/ml (low food concentration) of *Paramecium bursaria* for 48 h. After that, ca. 60 worms were pooled and washed with clean CM in preparation for RNA extraction. We performed three replicates for each feeding condition. RNA was extracted with the NucleoSpin RNA XS kit for RNA purification (Macherey-Nagel, 740902.50), according to the manufacturer’s standard protocol for RNA isolation from tissues, with the following modification: during the lysis step, samples were additionally snap-frozen in liquid nitrogen and thawed at room temperature to enhance tissue breakdown. The integrity of extracted RNA was evaluated on the Agilent 4150 TapeStation with the High Sensitivity RNA ScreenTape and sent for sequencing on an Illumina NovaSeq 6000 at the Genomics Core Facility (Centre of New Technologies, University of Warsaw) to a depth of 40 million 100-bp paired-end reads per sample.

Quality control of raw sequencing reads was assessed with FastQC (v0.12.1)^52^ and summarized with MultiQC (v1.18)^53^ before and after trimming. Adapter trimming was carried out using Trimmomatic (v0.39)^54^. Transcript abundances were quantified with Salmon (v1.10.2)^55^ against the *S*. *brevipharyngium* reference transcriptome (go_Sbre_v1; doi: 10.5281/zenodo.8239273)25. Transcript-level counts were imported into R (v4.3.3) using tximport (v1.30.0)^56^. Differential expression analysis was performed using DESeq2 (Bioconductor v3.18)^57^ with default settings. Log_2_ Fold Change (log_2_FC) was calculated as the ratio of low (27.5 cells/ml) versus high (275 cells/ml) food concentration. The Benjamini– Hochberg procedure was used to adjust p-values. Transcripts with adjusted *p* < 0.05 and |log_2_FC| > 2 were considered significantly differentially expressed. Principal component analysis (PCA) was performed on variance-stabilized expression values using the 500 most variable transcripts (Supplement Information 2). Open reading frames of differentially expressed transcripts were predicted and translated with TransDecoder (v5.7.1)^58^, using the –-single_best_only option. Functional annotation was generated using eggNOG-mapper (v2.1.13)^59^, employing DIAMOND (v2.1.23)^60^ for homology searches against the eggNOG database (v5.0.2)^61^. Volcano plots and row-scaled (z-score) heatmaps were produced for visualization. Processed transcriptomic datasets are provided in Supplement Information 2.

Cell-type-specific enrichment of differentially expressed genes was assessed in Python (v3.9.12) with Scanpy^62^, based on a published, annotated single-cell atlas of *S. brevipharyngium*^26^. For each gene and cell type, expression levels in cells belonging to a given cell type were compared to those in all remaining cells using a two-sided Wilcoxon rank-sum test. Enrichment was quantified as the fold change between mean expression within the cluster and mean expression across all other cells. P-values were adjusted for multiple testing using the Bonferroni correction. Genes were considered significantly enriched in a given cell type when the adjusted p-value was < 0.05 and fold change > 1. The number of enriched genes from the input gene set was then summarized for each cell type. Statistics for the single-cell analysis are available in Supplement Information 2.

A list of candidate marker genes was obtained from the single-cell transcriptomic data set of *S. brevipharyngium*^26^. Candidate genes were matched to the reference transcriptome using transcript identifiers. Differential expression statistics for candidate genes with adjusted *p* < 0.05 and |log_2_FC| > 1 were extracted from the DESeq2 results and visualized.

### Statistical analyses

All statistical analyses were performed in R (v4.5.1)^63^. In longitudinal data analyses of population dynamics under different food types, as well as chain formation and body length under different food concentrations, experimental day was treated as a continuous predictor and modeled using smooth terms^64,65^. Global differences were first tested by comparing nested models using chi-square tests. Final models were then used to estimate trajectories and to assess pairwise differences using Wald tests based on differences between model-predicted trajectories (Supplementary Tables 15, 16, and 17). For these analyses, as well as for time to first chain splitting, predator preference for prey size, and differences in the number of EdU+ cells, model fit was assessed using model-specific diagnostic procedures (Supplementary Tables 15, 16, 17, 18, 19, and 20). Pairwise comparisons were adjusted for multiple testing using the Holm method^66^.

Population dynamics (number of individuals) of *S. brevipharyngium*, *S. constrictum*, *S. simplex*, *S. bryophilum*, *S. leucops*, and *S. virginianum* were analyzed separately for each food type (*C. paramecium, C. variabilis, Cryptomonas sp., E. gracilis, L. inermis, P. bursaria, and S. obliquus*) using generalized additive models (GAMs) with a negative binomial distribution and a log link. Worm species was included as a fixed effect and replicate identity as a random effect (Supplementary Table 15). Additionally, reproductive rates (RR) were calculated as the average absolute change in population size per day,

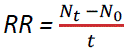

where *N*_0_ is the initial population size, *N*_t_ is the final population size, and *t* is the total observation time in days. Differences in RR among worm species within each food type were tested using Kruskal–Wallis tests followed by Dunn pairwise comparisons. Both analyses were used as complementary measures of population growth. Population dynamic analyses characterized temporal patterns of population growth throughout the experiment, whereas RR provided a summary measure of net population increase over the entire observation period^67^.

Changes in the number of zooids per chain under three food concentrations (275, 137.5, and 27.5 *P. bursaria* cells/ml) were analyzed separately for each worm species (*S. brevipharyngium, S. constrictum, S. simplex*, and *S. virginianum*) using generalized additive models (GAMs) with a Gamma distribution and log link. Food concentration was included as a fixed effect and replicate identity as a random effect (Supplementary Table 16). To assess whether differences in chain formation translated into differences in final population output, RR was also calculated using the number of individuals. Differences in RR among food concentrations within each worm species were tested using Kruskal–Wallis tests followed by Dunn pairwise comparisons.

*S. brevipharyngium* longitudinal growth rates under three food concentrations were analyzed using a generalized additive mixed model (GAMM) with a Gaussian distribution and identity link. Food concentration was included as a fixed effect and individual identity as a random effect (Supplementary Table 17).

Time to first chain splitting was analyzed using a Cox proportional hazards model^68^ with feeding regime (pulse vs. daily), three food concentrations, and their interaction as predictors. Time was measured as the day of the experiment, chain splitting was treated as the event of interest, and observations without splitting were treated as censored. Global effects of feeding regime, food concentration, and their interaction were evaluated using analysis of deviance with chi-square tests (Supplementary Table 18).

Predator preference for prey size within *S. brevipharyngium* or *S. virginianum* was analyzed using a generalized linear model (GLM) with a binomial error distribution and logit link. The model was fitted using cell-means parameterization to estimate the probability of selecting each prey size category for each prey species. Pairwise comparisons were obtained using the *emmeans* package (Supplementary Table 19)^69^.

Differences in the number of EdU+ cells adjusted for body length in *S. brevipharyngium* under three food concentrations were analyzed using a GLM with a Gaussian distribution and identity link. Food concentration was included as a fixed effect, and global effects were tested with an F-test. Pairwise differences among food concentrations were evaluated using estimated marginal means (Supplementary Table 20).

## Acknowledgements

We would like to thank Anna Karnkowska and Marta Sałek (Institute of Evolutionary Biology, University of Warsaw), as well as Anna Bednarska (Institute of Ecology, University of Warsaw), for sharing cultures of unicellular organisms used as food in our dietary experiments. The culture of *Chilomonas paramecium* originated from the Culture Collection of Algae of the University of Göttingen (SAG). We acknowledge the Laboratory of Imaging Tissue Structure and Function at the Nencki Institute, Leader of the Polish Euro-BioImaging Node “Advanced Light Microscopy NodePoland”, supported by the Minister of Education and Science based on contract No 2022/WK/05, for their assistance. Finally, we would like to thank Nicolas Bekkouche for his comments on the manuscript and figures.

## Funding

The research was funded by the Polish National Agency for Academic Exchange (Polish Returns NAWA grant no. BPN/PPO/2023/1/00002 to LG), National Science Centre, Poland (Polish Returns 2023 grant no. 2024/03/1/NZ8/00002 to LG), and Max-Planck-Gesellschaft (through Max Planck Partner Group funding to LG).

## Data availability

The raw experimental and microscopy data, as well as input datasets and scripts for statistical analyses, have been deposited at the Zenodo platform (https://doi.org/10.5281/zenodo.20488435). The raw RNA-seq reads of worms fed with different food concentrations have been deposited at the NCBI Sequence Reads Archive as BioProject PRJNA1499102.

## Authors contributions

LG designed the study, acquired funding, performed antibody staining and *in situ* hybridization, analyzed data, prepared figures, and drafted the manuscript. MS performed experiments (feeding, growth rate, EdU staining, predation), analyzed data statistically, cultured animals, prepared figures, and drafted parts of the manuscript. DK performed bioinformatic analyses, prepared figures, and drafted parts of the manuscript. KT performed feeding experiments, imaged animals on light and confocal microscopes, cultured animals, extracted RNA for transcriptome sequencing, prepared figures, and drafted parts of the manuscript.

**Extended Data Fig. 1.**
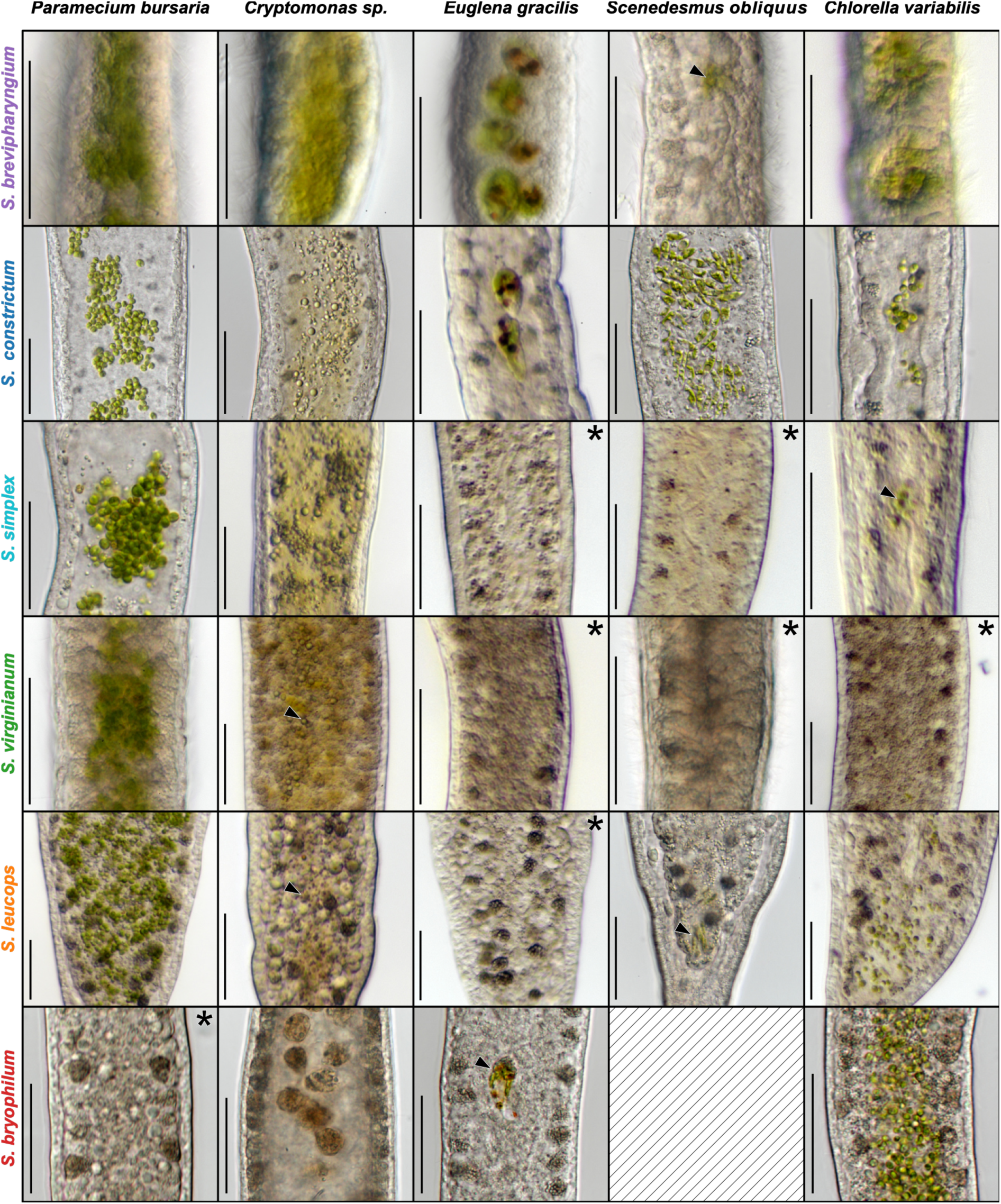
| Intake of *P. bursaria*, *Cryptomonas* sp., *E. gracilis*, *S. obliquus* and *C. variabilis* in different species of starved *Stenostomum* three hours after exposure to food. Asterisks indicate no intake of food. Scale bars represent 50 µm.

**Extended Data Fig. 2.**
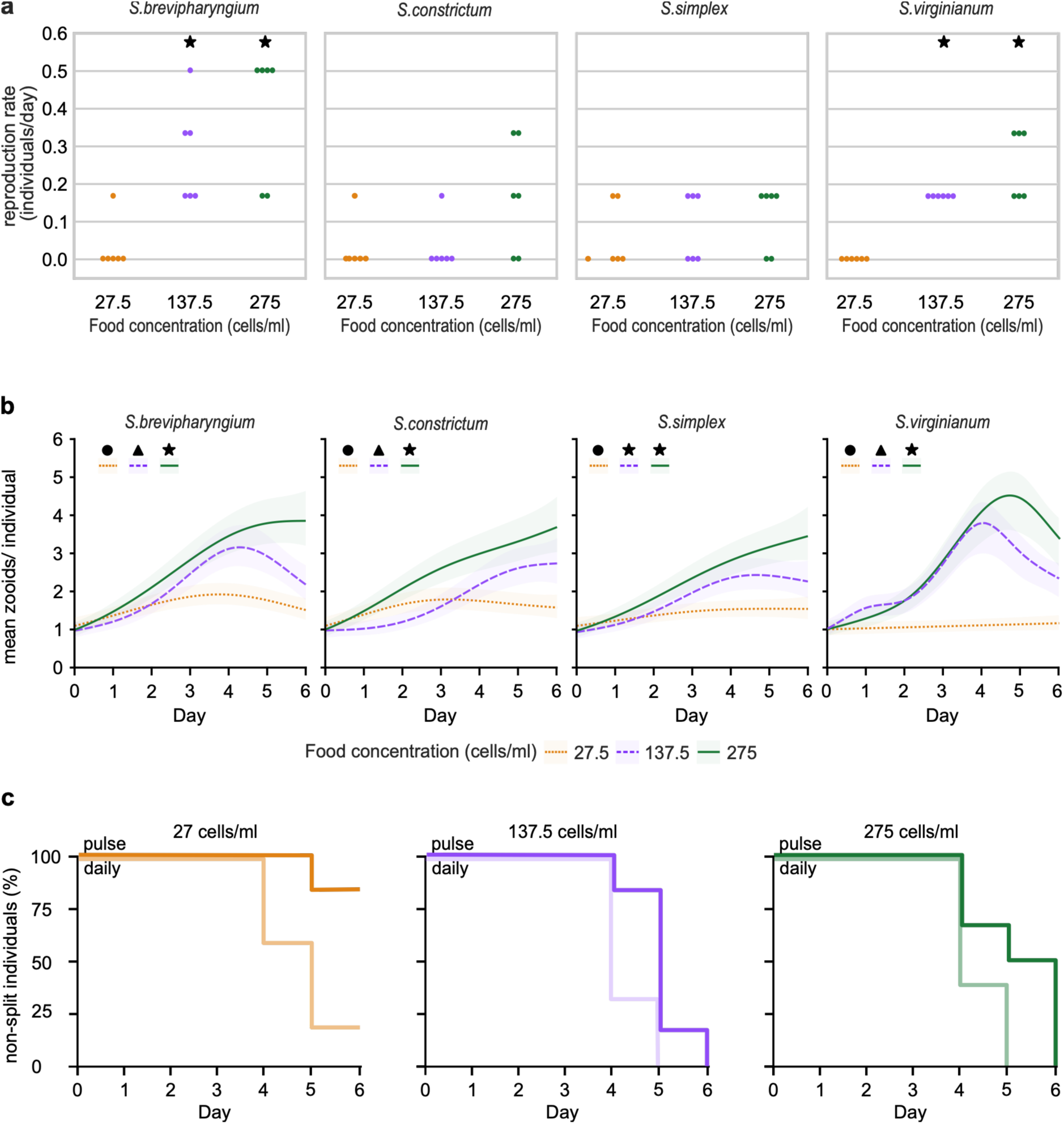
| Impact of the food concentration on worm development. **a** Asexual reproduction rates (RR; number of new individuals produced over 7 days) of individual founding worms in each experiment. Dots indicate RR values for single replicates. Stars indicate significant differences between food concentrations (p-value < 0.05). **B** Changes in the mean number of zooids per chain over one week for six *Stenostomum* species (N = 6 for each species–food concentration combination). Solid lines depict the changes in mean zooids per individual predicted by the generalized additive model (GAM), and lighter areas indicate the 95% confidence intervals. Different symbols (triangles, stars, and circles) indicate significant differences between trajectories. **c** Kaplan-Meier curves showing the percentage of non-split individuals of *S. brevipharyngium* over one week under two feeding regimes (pulse and daily) across three food concentrations. The lines represent survival trajectories, which are defined as the proportion of individuals that did not undergo splitting during the experiment. Cox proportional hazards analysis showed significant effects of feeding regime and food concentration, but not their interaction.

## Notes

### Competing Interest Statement

The authors have declared no competing interest.

