## Supplementary information 1 for "Excessive feeding induces development of a colony-like body plan in paratomous flatworms"

Table of Contents:

3: Figure S1: page 3

4–28: Tables S1–S20

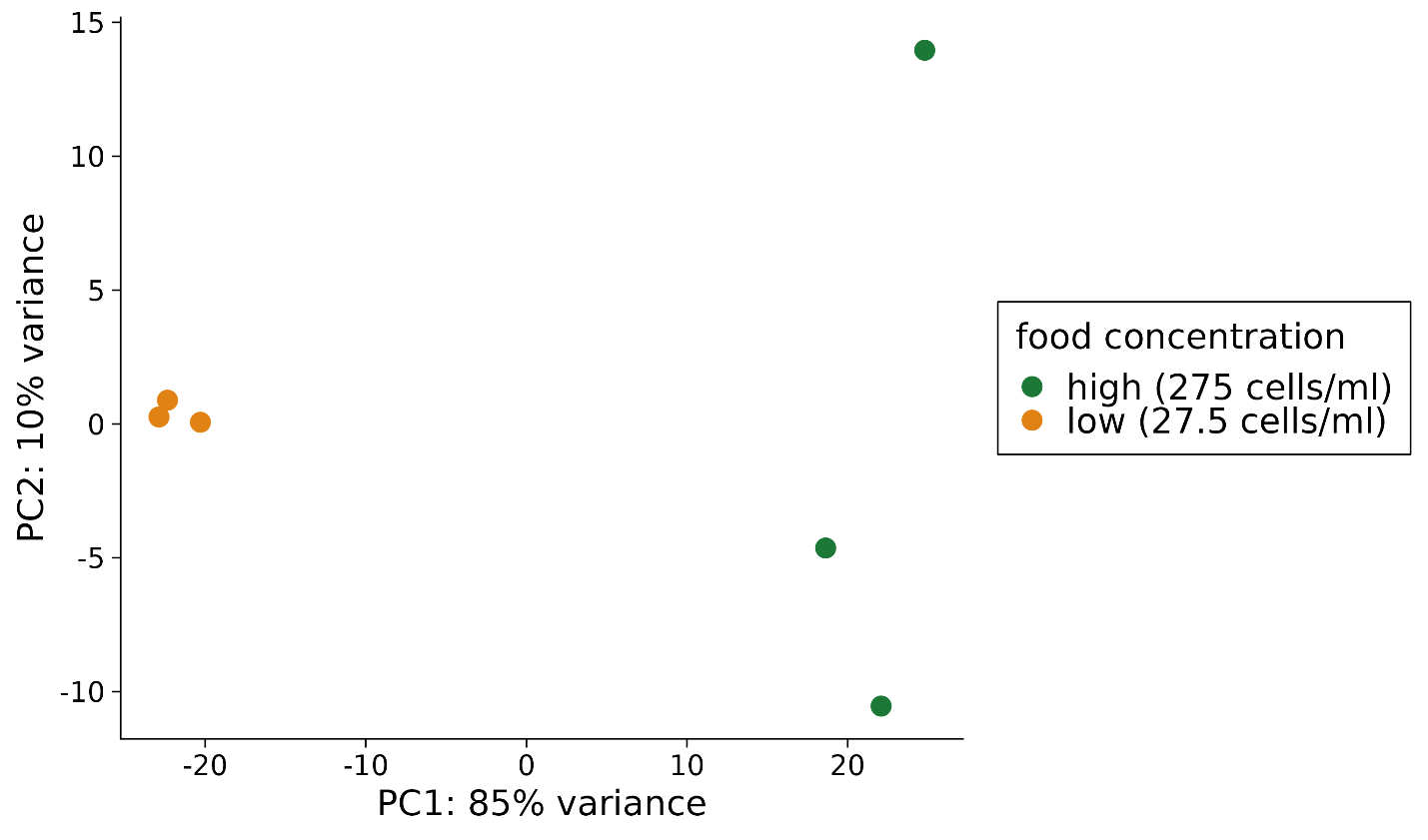
 **Supplementary Figure 1 | Principal component analysis (PCA) of variance-stabilized transcript abundance profiles in response to food availability.** Points represent individual biological replicates. Samples cultured under high food concentration (275 cells/ml; green) and low food concentration (27.5 cells/ml; orange) form distinct clusters.

**Supplementary Table 1 |** Global comparisons of population dynamics (number of individuals) of *S. brevipharyngium, S. constrictum, S. simplex, S. bryophyllum, S. leucops, and S. virginianum* under different food types: *C. paramecium, C. variabilis, Cryptomonas sp., E. gracilis, L. inermis, P. bursaria,* and *S. obliquus*, based on chi-square tests comparing nested generalized additive models (GAMs). The reduced model assumed a common temporal trajectory across species, whereas the full model allowed species-specific temporal trajectories. Significant p-values indicate that the full model differed significantly from the reduced model.

| **Food** | **Difference in df** | **Reduction in deviance** | **p-value** |
| --- | --- | --- | --- |
| *P. bursaria* | 1.5 | 31.8 | **p < 0.001** |
| *Cryptomonas* sp. | 5.00 | 88.0 | **p < 0.001** |
| *C. paramecium* | 5.00 | 21.6 | **p < 0.001** |
| *E. gracilis* | 6.52 | 119.0 | **p < 0.001** |
| *L. inermis* | 5.00 | 0.87 | p = 0.972 |
| *C. variabilis* | 5.00 | 7.27 | p = 0.201 |
| *S. obliquus* | 4.00 | 1.42 | p = 0.841 |

**Supplementary Table 2 |** Global comparisons of reproductive rates of *S. brevipharyngium, S. constrictum, S. simplex, S. bryophyllum, S. leucops,* and *S. virginianum* under different food types: *C. paramecium, C. variabilis, Cryptomonas* sp*., E. gracilis, L. inermis, P. bursaria,* and *S. obliquus*, based on the Kruskal–Wallis rank-sum test with a separate model fitted for each food type.

| **Food** | **χ²** | **df** | **p-value** |
| --- | --- | --- | --- |
| *P. bursaria* | 19.133 | 5 | **p = 0.002** |
| *Cryptomonas* sp. | 18.409 | 5 | **p = 0.002** |
| *C. paramecium* | 18.012 | 5 | **p = 0.003** |
| *E. gracilis* | 20.408 | 5 | **p = 0.001** |
| *L. inermis* | 4.182 | 5 | p = 0.5424 |
| *C. variabilis* | 12.504 | 5 | **p = 0.028** |
| *S. obliquus* | 6.517 | 4 | p = 0.164 |

**Supplementary Table 3 |** Pairwise comparisons of population dynamics (number of individuals) of *S. brevipharyngium*, *S. constrictum*, *S. simplex*, S*. bryophyllum*, *S. leucops*, *S. virginianum* under different food types: *C. paramecium*, *Cryptomonas sp*., *E. gracilis*, *P. bursaria*, based on Wald-type tests with Holm-adjusted p-values. *L. inermis, C. variabilis,* and *S. obliquus* are not shown because the global chi-square test comparing nested GAMs was not significant.

| **Food** | **Contrast** | **χ²** | **df** | **p-value** |
| --- | --- | --- | --- | --- |
| *P. bursaria* | *S. brevipharyngium * S. bryophyllum* | 34.4 | 13 | **0.011** |
| *P. bursaria* | *S. brevipharyngium * S. constrictum* | 1.6 | 13 | 1.000 |
| *P. bursaria* | *S. brevipharyngium * S. leucops* | 35.1 | 13 | **0.009** |
| *P. bursaria* | *S. brevipharyngium * S. simplex* | 4.23 | 13 | 1.000 |
| *P. bursaria* | *S. brevipharyngium * S. virginianum* | 2.37 | 13 | 1.000 |
| *P. bursaria* | *S. bryophyllum * S. constrictum* | 31.7 | 13 | **0.022** |
| *P. bursaria* | *S. bryophyllum * S. leucops* | 0.095 | 13 | 1.000 |
| *P. bursaria* | *S. bryophyllum * S. simplex* | 53.8 | 13 | **<0.001** |
| *P. bursaria* | *S. bryophyllum * S. virginianum* | 48.3 | 13 | **<0.001** |
| *P. bursaria* | *S. constrictum * S. leucops* | 32.5 | 13 | **0.018** |
| *P. bursaria* | *S. constrictum * S. simplex* | 7.65 | 13 | 1.000 |
| *P. bursaria* | *S. constrictum * S. virginianum* | 5.39 | 13 | 1.000 |
| *P. bursaria* | *S. leucops * S. simplex* | 54.4 | 13 | **<0.001** |
| *P. bursaria* | *S. leucops * S. virginianum* | 48.7 | 13 | **<0.001** |
| *P. bursaria* | *S. simplex * S. virginianum* | 0.601 | 13 | 1.000 |
| *Cryptomonas sp.* | *S. brevipharyngium * S. bryophyllum* | 158.0 | 13 | **<0.001** |
| *Cryptomonas sp.* | *S. brevipharyngium * S. constrictum* | 0.617 | 13 | 1.000 |
| *Cryptomonas sp.* | *S. brevipharyngium * S. leucops* | 3.800 | 13 | 1.000 |
| *Cryptomonas sp.* | *S. brevipharyngium * S. simplex* | 12.3 | 13 | 1.000 |
| *Cryptomonas sp.* | *S. brevipharyngium * S. virginianum* | 10.2 | 13 | 1.000 |
| *Cryptomonas sp.* | *S. bryophyllum * S. constrictum* | 163.0 | 13 | **<0.001** |
| *Cryptomonas sp.* | *S. bryophyllum * S. leucops* | 174.0 | 13 | **<0.001** |
| *Cryptomonas sp.* | *S. bryophyllum * S. simplex* | 177.0 | 13 | **<0.001** |
| *Cryptomonas sp.* | *S. bryophyllum * S. virginianum* | 178.0 | 13 | **<0.001** |
| *Cryptomonas sp.* | *S. constrictum * S. leucops* | 1.82 | 13 | 1.000 |
| *Cryptomonas sp.* | *S. constrictum * S. simplex* | 9.26 | 13 | 1.000 |
| *Cryptomonas sp.* | *S. constrictum * S. virginianum* | 7.37 | 13 | 1.000 |
| *Cryptomonas sp.* | *S. leucops * S. simplex* | 3.19 | 13 | 1.000 |
| *Cryptomonas sp.* | *S. leucops * S. virginianum* | 2.05 | 13 | 1.000 |
| *Cryptomonas sp.* | *S. simplex * S. virginianum* | 0.137 | 13 | 1.000 |
| *C. paramecium* | *S. brevipharyngium * S. bryophyllum* | 17.9 | 13 | 1.000 |
| *C. paramecium* | *S. brevipharyngium * S. constrictum* | 5.32 | 13 | 1.000 |
| *C. paramecium* | *S. brevipharyngium * S. leucops* | 17.9 | 13 | 1.000 |
| *C. paramecium* | *S. brevipharyngium * S. simplex* | 0.584 | 13 | 1.000 |
| *C. paramecium* | *S. brevipharyngium * S. virginianum* | 0.744 | 13 | 1.000 |
| *C. paramecium* | *S. bryophyllum * S. constrictum* | 36.3 | 13 | **0.008** |
| *C. paramecium* | *S. bryophyllum * S. leucops* | 0.000 | 13 | 1.000 |
| *C. paramecium* | *S. bryophyllum * S. simplex* | 23.0 | 13 | 0.458 |
| *C. paramecium* | *S. bryophyllum * S. virginianum* | 24.2 | 13 | 0.386 |
| *C. paramecium* | *S. constrictum * S. leucops* | 36.3 | 13 | **0.008** |
| *C. paramecium* | *S. constrictum * S. simplex* | 3.39 | 13 | 1.000 |
| *C. paramecium* | *S. constrictum * S. virginianum* | 2.12 | 13 | 1.000 |
| *C. paramecium* | *S. leucops * S. simplex* | 23.0 | 13 | 0.458 |
| *C. paramecium* | *S. leucops * S. virginianum* | 24.2 | 13 | 0.386 |
| *C. paramecium* | *S. simplex * S. virginianum* | 0.377 | 13 | 1.000 |
| *E. gracilis* | *S. brevipharyngium * S. bryophyllum* | 86.7 | 13 | **<0.001** |
| *E. gracilis* | *S. brevipharyngium * S. constrictum* | 10.3 | 13 | 1.000 |
| *E. gracilis* | *S. brevipharyngium * S. leucops* | 9.620 | 13 | 1.000 |
| *E. gracilis* | *S. brevipharyngium * S. simplex* | 0.115 | 13 | 1.000 |
| *E. gracilis* | *S. brevipharyngium * S. virginianum* | 3.08 | 13 | 1.000 |
| *E. gracilis* | *S. bryophyllum * S. constrictum* | 56.9 | 13 | **<0.001** |
| *E. gracilis* | *S. bryophyllum * S. leucops* | 31.1 | 13 | **0.036** |
| *E. gracilis* | *S. bryophyllum * S. simplex* | 87.9 | 13 | **<0.001** |
| *E. gracilis* | *S. bryophyllum * S. virginianum* | 84.4 | 13 | **<0.001** |
| *E. gracilis* | *S. constrictum * S. leucops* | 4.350 | 13 | 1.000 |
| *E. gracilis* | *S. constrictum * S. simplex* | 9.380 | 13 | 1.000 |
| *E. gracilis* | *S. constrictum * S. virginianum* | 7.79 | 13 | 1.000 |
| *E. gracilis* | *S. leucops * S. simplex* | 9.120 | 13 | 1.000 |
| *E. gracilis* | *S. leucops * S. virginianum* | 9.57 | 13 | 1.000 |
| *E. gracilis* | *S. simplex * S. virginianum* | 2.45 | 13 | 1.000 |

**Supplementary Table 4 |** Pairwise comparisons of reproductive rates of *S. brevipharyngium*, *S. constrictum*, *S. simplex*, S*. bryophyllum*, *S. leucops*, *S. virginianum* under different food types: *C. paramecium*, *C. variabilis*, *Cryptomonas sp*., *E. gracilis*, *P. bursaria,* based on Dunn tests with Holm-adjusted p-values. *L. inermis* and *S. obliquus* are not shown because the global Kruskal–Wallis test was not significant.

| **Food** | **Contrast** | **Z** | **p-value** |
| --- | --- | --- | --- |
| *P. bursaria* | *S. brevipharyngium * S. bryophyllum* | 2.063 | 0.431 |
| *P. bursaria* | *S. brevipharyngium * S. constrictum* | 0.433 | 1.000 |
| *P. bursaria* | *S. brevipharyngium * S. leucops* | 2.063 | 0.392 |
| *P. bursaria* | *S. brevipharyngium * S. simplex* | 1.171 | 1.000 |
| *P. bursaria* | *S. brevipharyngium * S. virginianum* | 0.789 | 1.000 |
| *P. bursaria* | *S. bryophyllum * S. constrictum* | 1.630 | 0.929 |
| *P. bursaria* | *S. bryophyllum * S. leucops* | 0.000 | 1.000 |
| *P. bursaria* | *S. bryophyllum * S. simplex* | 3.234 | **0.018** |
| *P. bursaria* | *S. bryophyllum * S. virginianum* | 2.852 | 0.057 |
| *P. bursaria* | *S. constrictum * S. leucops* | 1.630 | 0.825 |
| *P. bursaria* | *S. constrictum * S. simplex* | 1.604 | 0.761 |
| *P. bursaria* | *S. constrictum * S. virginianum* | 1.222 | 1.000 |
| *P. bursaria* | *S. leucops * S. simplex* | 3.234 | **0.017** |
| *P. bursaria* | *S. leucops * S. virginianum* | 2.852 | **0.052** |
| *P. bursaria* | *S. simplex * S. virginianum* | 0.382 | 1.000 |
| *Cryptomonas* sp. | *S. brevipharyngium * S. bryophyllum* | 3.442 | **0.009** |
| *Cryptomonas* sp. | *S. brevipharyngium * S. constrictum* | 0.000 | 1.000 |
| *Cryptomonas* sp. | *S. brevipharyngium * S. leucops* | 0.730 | 1.000 |
| *Cryptomonas* sp. | *S. brevipharyngium * S. simplex* | 1.982 | 0.570 |
| *Cryptomonas* sp. | *S. brevipharyngium * S. virginianum* | 1.982 | 0.475 |
| *Cryptomonas* sp*.* | *S. bryophyllum * S. constrictum* | 3.442 | **0.008** |
| *Cryptomonas* sp. | *S. bryophyllum * S. leucops* | 2.712 | 0.087 |
| *Cryptomonas* sp. | *S. bryophyllum * S. simplex* | 1.460 | 1.000 |
| *Cryptomonas* sp. | *S. bryophyllum * S. virginianum* | 1.460 | 1.000 |
| *Cryptomonas* sp*.* | *S. constrictum * S. leucops* | 0.730 | 1.000 |
| *Cryptomonas* sp. | *S. constrictum * S. simplex* | 1.982 | 0.523 |
| *Cryptomonas* sp. | *S. constrictum * S. virginianum* | 1.982 | 0.428 |
| *Cryptomonas* sp*.* | *S. leucops * S. simplex* | 1.252 | 1.000 |
| *Cryptomonas* sp. | *S. leucops * S. virginianum* | 1.252 | 1.000 |
| *Cryptomonas* sp. | *S. simplex * S. virginianum* | 0.000 | 1.000 |
| *C. paramecium* | *S. brevipharyngium * S. bryophyllum* | 2.383 | 0.224 |
| *C. paramecium* | *S. brevipharyngium * S. constrictum* | 0.785 | 1.000 |
| *C. paramecium* | *S. brevipharyngium * S. leucops* | 2.383 | 0.206 |
| *C. paramecium* | *S. brevipharyngium * S. simplex* | 0.131 | 1.000 |
| *C. paramecium* | *S. brevipharyngium * S. virginianum* | 0.131 | 1.000 |
| *C. paramecium* | *S. bryophyllum * S. constrictum* | 3.168 | **0.023** |
| *C. paramecium* | *S. bryophyllum * S. leucops* | 0.000 | 1.000 |
| *C. paramecium* | *S. bryophyllum * S. simplex* | 2.252 | 0.268 |
| *C. paramecium* | *S. bryophyllum * S. virginianum* | 2.252 | 0.219 |
| *C. paramecium* | *S. constrictum * S. leucops* | 3.168 | **0.022** |
| *C. paramecium* | *S. constrictum * S. simplex* | 0.916 | 1.000 |
| *C. paramecium* | *S. constrictum * S. virginianum* | 0.916 | 1.000 |
| *C. paramecium* | *S. leucops * S. simplex* | 2.252 | 0.243 |
| *C. paramecium* | *S. leucops * S. virginianum* | 2.252 | 0.195 |
| *C. paramecium* | *S. simplex * S. virginianum* | 0.000 | 1.000 |
| *E. gracilis* | *S. brevipharyngium * S. bryophyllum* | 1.336 | 1.000 |
| *E. gracilis* | *S. brevipharyngium * S. constrictum* | 2.298 | 0.259 |
| *E. gracilis* | *S. brevipharyngium * S. leucops* | 2.298 | 0.237 |
| *E. gracilis* | *S. brevipharyngium * S. simplex* | 0.267 | 1.000 |
| *E. gracilis* | *S. brevipharyngium * S. virginianum* | 1.282 | 1.000 |
| *E. gracilis* | *S. bryophyllum * S. constrictum* | 3.633 | **0.004** |
| *E. gracilis* | *S. bryophyllum * S. leucops* | 3.633 | **0.004** |
| *E. gracilis* | *S. bryophyllum * S. simplex* | 1.603 | 0.872 |
| *E. gracilis* | *S. bryophyllum * S. virginianum* | 2.618 | 0.115 |
| *E. gracilis* | *S. constrictum * S. leucops* | 0.000 | 1.000 |
| *E. gracilis* | *S. constrictum * S. simplex* | 2.030 | 0.423 |
| *E. gracilis* | *S. constrictum * S. virginianum* | 1.015 | 1.000 |
| *E. gracilis* | *S. leucops * S. simplex* | 2.030 | 0.381 |
| *E. gracilis* | *S. leucops * S. virginianum* | 1.015 | 1.000 |
| *E. gracilis* | *S. simplex * S. virginianum* | 1.015 | 0.930 |
| *C. variabilis* | *S. brevipharyngium * S. bryophyllum* | 2.914 | 0.054 |
| *C. variabilis* | *S. brevipharyngium * S. constrictum* | 2.285 | 0.313 |
| *C. variabilis* | *S. brevipharyngium * S. leucops* | 0.762 | 1.000 |
| *C. variabilis* | *S. brevipharyngium * S. simplex* | 2.285 | 0.290 |
| *C. variabilis* | *S. brevipharyngium * S. virginianum* | 2.285 | 0.268 |
| *C. variabilis* | *S. bryophyllum * S. constrictum* | 0.629 | 1.000 |
| *C. variabilis* | *S. bryophyllum * S. leucops* | 2.152 | 0.345 |
| *C. variabilis* | *S. bryophyllum * S. simplex* | 0.629 | 1.000 |
| *C. variabilis* | *S. bryophyllum * S. virginianum* | 0.629 | 1.000 |
| *C. variabilis* | *S. constrictum * S. leucops* | 1.523 | 1.000 |
| *C. variabilis* | *S. constrictum * S. simplex* | 0.000 | 1.000 |
| *C. variabilis* | *S. constrictum * S. virginianum* | 0.000 | 1.000 |
| *C. variabilis* | *S. leucops * S. simplex* | 1.523 | 1.000 |
| *C. variabilis* | *S. leucops * S. virginianum* | 1.523 | 1.000 |
| *C. variabilis* | *S. simplex * S. virginianum* | 0.000 | 1.000 |

**Supplementary Table 5 |** Global comparisons of reproductive rates in *S. brevipharyngium, S. constrictum, S. simplex,* and *S. virginianum* under different food concentrations: 27.5, 137.5, and 275 *P. bursaria* cells ml⁻¹, based on the Kruskal–Wallis rank-sum test, with a separate analysis conducted for each species.

| **Worm species** | **χ²** | **df** | **p-value** |
| --- | --- | --- | --- |
| *S. brevipharyngium* | 11.349 | 2 | **p = 0.003** |
| *S. constrictum* | 5.06 | 2 | p = 0.08 |
| *S. simplex* | 1.259 | 2 | p = 0.533 |
| *S. virginianum* | 14.733 | 2 | **p < 0.001** |

**Supplementary Table 6 |** Pairwise comparisons of reproductive rates of *S. brevipharyngium* and *S. virginianum* under different food concentrations: 27.5, 137.5, and 275 *P. bursaria* cells ml⁻¹, based on Dunn tests with Holm-adjusted p-values. *S. constrictum* and *S. simplex* are not shown because the global Kruskal–Wallis test was not significant.

| **Worm species** | **Contrast** | **Z** | **p-value** |
| --- | --- | --- | --- |
| *S. brevipharyngium* | 27.5 * 137.5 cells ml⁻¹ | -2.422 | **0.03** |
| *S. brevipharyngium* | 27.5 * 275 cells ml⁻¹ | 3.239 | **0.003** |
| *S. brevipharyngium* | 275 * 137.5 cells ml⁻¹ | 0.817 | 0.414 |
| *S. virginianum* | 27.5 * 137.5 cells ml⁻¹ | -2.661 | **0.015** |
| *S. virginianum* | 27.5 * 275 cells ml⁻¹ | 3.726 | **<0.001** |
| *S. virginianum* | 275 * 137.5 cells ml⁻¹ | 1.064 | 0.287 |

**Supplementary Table 7 |** Global comparisons of changes in the number of zooids per chain in *S. brevipharyngium, S. constrictum, S. simplex, S. virginianum* under three food concentrations: 27.5, 137.5, and 275 *P. bursaria* cells ml⁻¹, based on chi-square tests comparing nested generalized additive models (GAMs). The reduced model assumed a common temporal trajectory across food concentrations, whereas the full model allowed food-specific temporal trajectories. Significant p-values indicate that the full model differed significantly from the reduced model.

| **Worm species** | **Difference in df** | **Reduction in deviance** | **p-value** |
| --- | --- | --- | --- |
| *S. brevipharyngium* | 9.66 | 4.03 | **p < 0.001** |
| *S. constrictum* | 6.99 | 4.05 | **p < 0.001** |
| *S. simplex* | 12.16 | 3.18 | **p < 0.001** |
| *S. virginianum* | 10.27 | 6.39 | **p < 0.001** |

**Supplementary Table 8 |** Pairwise comparisons of changes in the number of zooids per chain in *S. brevipharyngium, S. constrictum, S. simplex, S. virginianum* under three food concentrations: 27.5, 137.5, and 275 *P. bursaria* cells ml⁻¹, based on Wald-type tests with Holm-adjusted p-values.

| **Worm species** | **Contrast** | **χ²** | **df** | **p-value** |
| --- | --- | --- | --- | --- |
| *S. brevipharyngium* | 27.5 * 137.5 cells ml⁻¹ | 32.0 | 7 | **p < 0.001** |
| *S. brevipharyngium* | 27.5 * 275 cells ml⁻¹ | 64.1 | 7 | **p < 0.001** |
| *S. brevipharyngium* | 275 * 137.5 cells ml⁻¹ | 19.3 | 7 | **p = 0.007** |
| *S. constrictum* | 27.5 * 137.5 cells ml⁻¹ | 49.1 | 7 | **p < 0.001** |
| *S. constrictum* | 27.5 * 275 cells ml⁻¹ | 51.6 | 7 | **p < 0.001** |
| *S. constrictum* | 275 * 137.5 cells ml⁻¹ | 25.9 | 7 | **p < 0.001** |
| *S. simplex* | 27.5 * 137.5 cells ml⁻¹ | 23.7 | 7 | **p = 0.003** |
| *S. simplex* | 27.5 * 275 cells ml⁻¹ | 52.0 | 7 | **p < 0.001** |
| *S. simplex* | 275 * 137.5 cells ml⁻¹ | 10.7 | 7 | p = 0.154 |
| *S. virginianum* | 27.5 * 137.5 cells ml⁻¹ | 156.0 | 7 | **p < 0.001** |
| *S. virginianum* | 27.5 * 275 cells ml⁻¹ | 218.0 | 7 | **p < 0.001** |
| *S. virginianum* | 275 * 137.5 cells ml⁻¹ | 25.0 | 7 | **p < 0.001** |

**Supplementary Table 9 |** Global effects of feeding regime (pulse and daily), food concentration (27.5, 137.5, and 275 *P. bursaria* cells ml⁻¹), and their interaction on time to first chain splitting in *S. brevipharyngium*, based on analysis of deviance of the Cox proportional hazards model.

| **Predictor** | **χ²** | **df** | **p-value** |
| --- | --- | --- | --- |
| Feeding regime | 7.322 | 1 | **0.007** |
| Food concentration | 11.894 | 2 | **0.003** |
| Feeding regime × Food concentration | 2.093 | 2 | 0.351 |

**Supplementary Table 10 |** Pairwise comparisons of prey size within each prey species preferred by the predator. Results are based on estimated marginal means from the binomial GLM, with z-tests adjusted for multiple comparisons using the Holm method.

| **Prey species** | **Contrast** | **Odds ratio** | **Standard error** | **z-value** | **p-value** |
| --- | --- | --- | --- | --- | --- |
| *S. brevipharyngium* | 1-zooid * 3-zooid | 0.012 | 0.018 | -2.948 | **0.003** |
| *S. virginianum* | 1-zooid * 3-zooid | 0.25 | 0.25 | -1.386 | 0.166 |

**Supplementary Table 11 |** Global comparison of *S. brevipharyngium* longitudinal growth rate under three food concentrations: 27.5, 137.5, and 275 *P. bursaria* cells ml⁻¹, based on a chi-square test comparing nested generalized additive mixed models (GAMMs). The reduced model assumed a common temporal trajectory across treatments, whereas the full model allowed treatment-specific temporal trajectories. Significant p-values indicate that the full model differed significantly from the reduced model.

| **Difference in df** | **Reduction in deviance** | **p-value** |
| --- | --- | --- |
| 11.482 | 1.877 | **<0.0001** |

**Supplementary Table 12 |** Pairwise comparisons of *S. brevipharyngium* longitudinal growth rate among three food concentrations (27.5, 137.5, and 275 cells ml⁻¹), based on Wald-type tests with Holm-adjusted p-values.

| **Contrast** | **χ²** | **df** | **p-value** |
| --- | --- | --- | --- |
| 27.5 * 137.5 cells ml⁻¹ | 288.708 | 5 | **<0.0001** |
| 27.5 * 275 cells ml⁻¹ | 153.371 | 5 | **<0.0001** |
| 275 * 137.5 cells ml⁻¹ | 10.353 | 5 | 0.0658 |

**Supplementary Table 13 |** Global effect of food concentration on the number of EdU+ cells adjusted for body length in *S. brevipharyngium*, assessed in a Gaussian generalized linear model (GLM) using analysis of deviance with an F-test.

| **Effect** | **df** | **F** | **p-value** |
| --- | --- | --- | --- |
| Food concentration | 2 | 20.332 | **<0.001** |

**Supplementary Table 14 |** Pairwise comparisons of the number of EdU+ cells adjusted for body length in *S. brevipharyngium* among three food concentrations (27.5, 137.5, and 275 *P. bursaria* cells ml⁻¹). Results are based on estimated marginal means from the Gaussian GLM, with Holm-adjusted p-values.

| **Contrast** | **Estimate** | **Standard error** | **df** | **p-value** |
| --- | --- | --- | --- | --- |
| 27.5 vs 137.5 cells ml⁻¹ | -0.489 | 0.121 | 9 | **0.006** |
| 27.5 vs 275 cells ml⁻¹ | -0.763 | 0.121 | 9 | **<0.001** |
| 137.5 vs 275 cells ml⁻¹ | -0.275 | 0.121 | 9 | **0.05** |

**Supplementary Table 15 |** Structure of the generalized additive model (GAM) used to analyze population dynamics (number of individuals) of *S. brevipharyngium*, *S. constrictum*, *S. simplex*, S*. bryophyllum*, *S. leucops*, *S. virginianum* under different food types: *C. paramecium*, *C. variabilis*, *Cryptomonas sp*., *E. gracilis*, *L. inermis*, *P. bursaria*, *S. obliquus*. Separate models were fitted for each food type.

| **Model component** | **Specification** |
| --- | --- |
| Distribution / link | Negative binomial / log |
| Response variable | Population size (number of individuals) |
| Fixed and smooth effects | Worm species + smooth effect of day + species-specific smooth deviations |
| Random effect | Replicate identity |
| Model diagnostic procedures | *gam.check()*; Pearson residuals vs fitted values; QQ-plots and histograms of deviance residuals; observed vs fitted plots |
| Number of replicate trajectories | 24 per food* |
| Observations  * Except *S. obliquus* = 260  ** Except *S. obliquus* = 20 | 312 per food type** |

**Supplementary Table 16 |** Structure of the generalized additive model (GAM) used to analyze changes in the number of zooids per chain in *S. brevipharyngium*, S*. constrictum*, *S. simplex*, *S. virginianum* under three food concentrations: 27.5, 137.5, and 275 *P. bursaria* cells ml⁻¹. Separate models were fitted for each worm species.

| **Model component** | **Specification** |
| --- | --- |
| Distribution / link | Gamma / log |
| Response variable | Mean number of zooids per individual |
| Fixed and smooth effects | Food concentration + smooth effect of day + concentration-specific smooth deviations |
| Random effect | Replicate identity |
| Basis dimension of smooth term | k = 5 for most species; k = 7 for *S. virginianum* |
| Model diagnostic procedures | *gam.check()*; Pearson residuals vs fitted values; QQ-plots and histograms of deviance residuals; observed vs fitted plots |
| Number of replicate trajectories | 18 |
| Observations | 126 |

**Supplementary Table 17 |** Structure of the generalized additive mixed model (GAMM) used to analyze *S. brevipharyngium* longitudinal growth rate during exposure to three food concentrations: 27.5, 137.5, and 275 *P. bursaria* cells ml⁻¹.

| **Model component** | **Specification** |
| --- | --- |
| Distribution / link | Gaussian / identity |
| Response variable | Body length |
| Fixed and smooth effects | Food concentration + s(day) + food concentration-specific smooth deviations |
| Random effect | Individual ID |
| Correlation structure | AR (1) within individual |
| Variance structure | Food concentration-specific residual variance |
| Model diagnostic procedures | Normalized residuals vs fitted values; QQ-plots of normalized residuals |
| Number of individuals | 24 |
| Observations | 120 |

**Supplementary Table 18 |** Structure of the Cox proportional hazards model used to analyze the timing of first chain splitting in *S. brevipharyngium* fed with different concentrations of *P. bursaria* (27.5, 137.5, and 275 cells ml⁻¹) under pulse and daily feeding regimes.

| **Model component** | **Specification** |
| --- | --- |
| Model type | Cox proportional hazards model |
| Response variable | Time to first chain splitting |
| Fixed effects | *Feeding regime + food concentration + feeding regime × food concentration* |
| Censoring | Individuals without chain splitting during the observation |
| Model diagnostic procedures | Schoenfeld residuals; deviance residuals; DFBETA influence diagnostics |
| Observations | 34 |
| Number of first chain-splitting events | 28 |
| Censored observations | 6 |

**Supplementary Table 19** **|** Structure of the generalized linear model (GLM) used to analyze the effect of prey size (1-zooid vs 3-zooid) on prey choice by the predator *S. sphagnetorum* across two prey species (*S. brevipharyngium* and *S. virginianum*).

| **Model component** | **Specification** |
| --- | --- |
| Distribution / link | Binomial / logit |
| Response variable | Binary prey choice (chosen vs not chosen; 0/1) |
| Fixed effects | Prey size × Prey species |
| Model parameterization | Cell-means |
| Model diagnostic procedures | DHARMa diagnostics (residual distribution, dispersion, zero-inflation), separation check |
| Number of choice trials | 19 |
| Observations (prey items) | 38 |

**Supplementary Table 20 |** Structure of the generalized linear model (GLM) used to analyze the number of EdU+ cells adjusted for body length in *S. brevipharyngium* during exposure to three food concentrations: 27.5, 137.5, and 275 *P. bursaria* cells ml⁻¹.

| **Model component** | **Specification** |
| --- | --- |
| Distribution / link | Gaussian / identity |
| Response variable | EdU-positive cells / body length |
| Fixed effects | Food concentration |
| Model diagnostic procedures | Shapiro–Wilk test, Levene’s test, Breusch–Pagan test, QQ-plot, residuals vs fitted plot, standardized residuals |
| Observations | 12 |
